# CCS Predictor 2.0: An Open-Source Jupyter Notebook Tool for Filtering Out False Positives in Metabolomics

**DOI:** 10.1101/2022.08.09.503345

**Authors:** Markace A. Rainey, Chandler A. Watson, Carter K. Asef, Makayla R. Foster, Erin S. Baker, Facundo M. Fernández

## Abstract

Metabolite annotation continues to be the widely accepted bottleneck in non-targeted metabolomics workflows. Annotation of metabolites typically relies on a combination of high resolution mass spectrometry (MS) with parent and tandem measurements, isotope cluster evaluations, and Kendrick mass defect (KMD) analysis. Chromatographic retention time matching with standards is often used at the later stages of the process, which can also be followed by metabolite isolation and structure confirmation utilizing nuclear magnetic resonance (NMR) spectroscopy. The measurement of gas phase collision cross section (CCS) values by ion mobility (IM) spectrometry also adds an important dimension to this workflow by generating an additional molecular parameter that can be used for filtering unlikely structures. The millisecond timescale of IM spectrometry allows the rapid measurement of CCS values and allows easy pairing with existing MS workflows. Here, we report on a highly accurate machine learning algorithm (CCSP 2.0) in an open-source Jupyter Notebook format to predict CCS values based on linear support vector regression models. This tool allows customization of the training set to the needs of the user, enabling the production of models for new adducts or previously unexplored molecular classes. CCSP produces predictions with accuracy equal to or greater than existing machine learning approaches such as CCSbase, DeepCCS and AllCCS, while being better aligned with FAIR (Findable, Accessible, Interoperable and Reusable) data principles. Another unique aspect of CCSP 2.0 its inclusion of a large library of 1613 molecular descriptors via the Mordred Python package, further encoding the fine aspects of isomeric molecular structures. CCS prediction accuracy was tested using CCS values in the McLean CCS Compendium with median relative errors of 1.25, 1.73 and 1.87% for the 170 [M-H]^-^, 155 [M+H]^+^ and 138 [M+Na]^+^ adducts tested. For class-matched data sets, CCS predictions via CCSP allowed filtering of 36.1% of incorrect structures while retaining a total of 100% of the correct annotations using a ΔCCS threshold of 2.8% and a mass error of 10 ppm.

## INTRODUCTION

Metabolomics is the least developed “omics” field, focusing on the examination of metabolites in complex systems. It has the potential to catalog the vast chemical diversity arising from metabolic activity among the millions of unique organisms on Earth by mapping cellular pathways whose trajectories correlate with the onset and progression of specific processes, such as cell differentiation, organismal growth and development, and disease^1, 2^. The metabolome is a collection of small molecules in a biological system that fall roughly under the ∼1500 Da molecular weight window. These molecules include endogenous species that are biosynthesized in “primary metabolism”, specialized “secondary metabolite” signaling molecules, lifestyle or environmental exposure molecules (the exposome), and molecules originating from the microbial community associated with the organism under study (the microbiome).

One of the major limitations in metabolomics is the identification of unknown molecules. The primary metabolomics platforms presently used include nuclear magnetic resonance (NMR) spectroscopy and chromatographic separations coupled with mass spectrometry (MS), such as liquid chromatography-mass spectrometry (LC-MS) and gas chromatography-mass spectrometry (GC-MS).^3^ Currently, the most popular solutions to structure elucidation involve combinations of several MS approaches such as high resolution MS, tandem MS, and sequential tandem MS (MS^n^)^4^. Even with these tools, metabolite identification remains the undisputed bottleneck in metabolomics workflows, with fewer than 2-10% of the detected compounds in typical non-targeted metabolomics studies annotated reliably^5^. The exploration of the remaining chemical space, sometimes described as the metabolome “dark matter”^6^, continues to be the focus of intense scientific research.

Ion mobility (IM) spectrometry coupled to MS has emerged as a robust platform to aid in metabolite identification^7, 8^. IM can be thought of as analogous to a gas phase electrophoretic experiment, where ions in a buffer gas pass through a chamber in the presence of an electric field, and their migration or drift time is measured. This drift time can then be used to calculate a collision cross section (CCS) value through appropriate regression or calibration procedures^9^. IM CCS values, while partially correlated with mass-to-charge ratios (*m/z*), are also dependent on molecular shape, granting IM-MS instrumentation a good degree of orthogonality for distinguishing isomers and isobars^10^. Moreover, CCS values have higher inter-laboratory robustness than chromatographic retention times^11^ and are starting to be compiled in databases for unknown metabolite annotation^12^.

CCS values for molecules as small as metabolites and as large as protein complexes have been predicted computationally for comparison to IM measurements^13, 14^. However, such predictions require 3-dimensional structures by way of solid-state NMR or X-ray crystallography experimental data. Quantum methods have shown significant promise in prediction of CCS values but tend to be computationally expensive^15-17^. As a faster, less resource-intensive alternative, several efforts involving machine learning (ML) CCS prediction have been reported in the literature. These include support vector regression (SVR) models such as MetCCS, LipidCCS and AllCCS^18-20^, more sophisticated regression models such as CCSBase^20^, joint prediction of CCS and chromatographic retention times^21^, and deep neural network models such as DeepCCS^22^, among others. However, challenges include tailoring the approaches to the data sets of interest, transparency or customizability of the ML models, and/or whether high accuracy CCS values can be generated for the analytes in question.

Here, we describe CCS Predictor 2.0 (CCSP 2.0), an open-source tool in a Jupyter Notebook format for ease of use by both experienced IM scientists and novice trainees alike. The goal of this tool is to produce CCS value predictions with accuracy equal to or greater than existing ML approaches, while aligning with FAIR (Findable, Accessible, Interoperable and Reusable) data principles^23^. CCSP 2.0 is illustrated using 463 test CCS measurements retrieved from the McLean CCS Compendium^24^, resulting in relative median errors lower than 2% for sodium adducts and protonated and deprotonated molecules. While several tools exist for CCS value predictions, CCSP provides a new framework for users to customize their own CCS predictions based on their unique training sets and workflow needs.

## Methods

### CCSP 2.0 Package Dependencies

CCSP 2.0 was developed in JupyterLab V2.2.6 using Python V3.8.5 and the following dependencies: RDKit V2020.09.1, PubChemPy V1.0.4, Pandas V1.1.3, Numpy V1.19.2, Matplotlib V3.3.2, and Plotly V4.14.3. The full CCSP 2.0 Jupyter notebook is publicly available at https://github.com/facundof2016/CCSP2.0.

### Data Selection and Scrubbing

All drift tube collision cross section measurements utilized for model creation and evaluation were obtained from the McLean Unified CCS Compendium^24^. Complete data sets were downloaded for [M-H]^-^, [M+H]^+^, and [M+Na]^+^ ions in the form of separate Microsoft Excel spreadsheets containing all corresponding entries in the Compendium (Supplemental Tables 1-3). In cases where duplicate entries existed in the McLean database, these were replaced with a single entry using the average of the provided CCS values. Database entries were excluded if the mass-to-charge ratio (*m/z*) corresponding to the provided IUPAC International Chemical Identifier (InChI) and specific ionic species deviated from the listed theoretical *m/z* by more than 1 mDa, which typically resulted from the improper representation of isotopically-labeled compounds. Database entries were also excluded if they contained a chemical element found in fewer than five database entries. Additional database entries were excluded if they were not compatible with the mass ranges set in the ML algorithms used by DeepCCS, AllCCS, or CCSBase. Following these data cleaning steps, 567 singly deprotonated ions, 518 singly protonated ions and 461 singly sodiated ions remained.

### Matrix Representation of Molecules and Further Data Preparation

Each molecule in the McLean database was initially represented by its neutral InChI string, which was then converted to the corresponding isomeric simplified molecular-input line-entry system (SMILES) specification using the Identifier Exchange Service (PubChem service). RDKit was used to encode molecular data (MOL) files for each SMILES. Each MOL file was used to calculate 1613 2D molecular descriptors using the Python package Mordred V1.2.0. Each iteration of the ML algorithm randomly assigned 70% of the molecules in the data set to a calibration (training) group for building models and the remaining 30% were withheld for external validation (testing). Molecular descriptors that were constant across all calibration molecules were discarded and any descriptor not applicable to one or more molecules in the calibration or validation sets was excluded. The most common category of inapplicable descriptors attempted to find the average of a value across bond paths; smaller molecules often lack bond paths that satisfy the desired bond count, causing these molecular descriptors to return a division-by-zero error. The remaining descriptors were *Z*-transformed using the mean and standard deviation of each descriptor in the training set. Any descriptor with a test datapoint that exceeded 1000 standard deviations from its mean was excluded.

### Model Development and Evaluation

The three data subsets ([M-H]^-^, [M+H]^+^, and [M+Na]^+^) were next used to train linear kernel support vector regression (SVR) models. Two hyperparameters were tuned in model development: (1) the regularization parameter *C*, which determines the penalty gradient associated with non-zero residuals and (2) *ϵ*, the radius of an epsilon-tube in which no penalty is accrued for non-zero residuals. Hyperparameter tuning followed a grid search approach, where *C* and *ϵ* were evaluated pairwise using 5-fold cross-validation. *C* was selected from seven options (0.00390625, 0.0078125, 0.015625, 0.03125, 0.0625, 0.125, and 0.25), and *ϵ* from three options (0.5, 1, 5). Hyperparameters yielding the lowest root mean squared error of cross-validation (RMSECV) were selected for the construction of the initial SVR model. Linear SVR model construction and hyperparameter tuning were performed using the SVR and GridSearchCV modules of SciKit-Learn V0.23.2 (scikit-learn.org), respectively. The initial SVR model provided features weights that were used to rank the importance of each molecular descriptor. Recursive Feature Elimination (RFE) with 5-fold cross-validation was used to simplify the SVR models by reducing the number of molecular descriptors utilized. The SciKit-Learn RFECV module was used to iteratively remove the five lowest-ranked molecular descriptors until the RMSECV increased. A final SVR model was constructed by repeating the hyperparameter tuning process using only the RFE-selected descriptors. Model validation was performed by evaluating the calibration, cross-validation, and validation errors through the median relative error (MRE), root mean squared error (RMSE), and coefficient of correlation (R^2^).

### Comparison of CCS Prediction Tools

Each adduct data set was repeatedly sampled, selecting 30% of a data set in each iteration for comparison between AllCCS, CCSbase, DeepCCS and CCSP 2.0. The same isomeric SMILES specifications were used as inputs for each tool, and the predicted CCS values of the metabolites from each tool were compared to the experimentally-measured values in the McLean Compendium. AllCCS and CCSbase predictions were completed with the corresponding web server, while DeepCCS predictions were performed using its built-in command line model. CCSP .0 was trained with the remaining 70% of each data set. All three adduct types were tested 101 times to provide an estimate of sampling variability in terms of MRE and RMSE.

### Evaluation of CCS Filtering Efficacy for Chemically Diverse Data Sets

To evaluate the efficacy of filtering metabolite annotation candidates *via* CCS predictions based on diverse training sets, a single 70:30 split of the [M-H]^-^ data set was utilized (Supplemental Table 4). A total of 397 compounds were allocated to model calibration, while 170 compounds were utilized for model validation and filtering assessment. For each compound in the validation set, a list of candidate annotations was obtained by querying PubChem using the Entrez Global Query Cross-Database Search System. The annotation candidate list for each standard consisted of all PubChem compounds with an exact mass within 10 parts-per-million (ppm) of the standard’s exact mass as calculated by Mordred. The querying process was automated using the Entrez module of the Biopython project, where the results listed each candidate using its PubChem Chemical Identifier (CID). The PubChem file for each CID was obtained using the Python module PubChemPy V1.0.4, and the corresponding isomeric SMILES was used to calculate their molecular descriptors. Nine of the validation compounds did not return exact PubChem matches and were excluded from filtering analysis. The remaining candidate sets were then used for filtering evaluation and receiver operating characteristic curve (ROC) analysis.

### Evaluation of CCS Filtering Efficacy for Class-Matched Data Sets

To evaluate the efficacy of filtering metabolite annotation candidates *via* CCS predictions based on a class-matched training set, a single 70:30 split of the [M-H]^-^ lipid and lipid-like molecular superclass was utilized (Supplemental Table 5). A total of 93 compounds were allocated to model calibration, while 36 compounds were utilized for model validation and filtering assessment. For each compound in the validation set, a list of candidate annotations was obtained by querying PubChem using the Entrez Global Query Cross-Database Search System. The annotation candidate list for each standard consisted of all PubChem compounds with an exact mass within 10 parts-per-million (ppm) of the standard’s exact mass as calculated by Mordred. The querying process was automated using the Entrez module of the Biopython project, where the results listed each candidate using its PubChem Chemical Identifier (CID). The PubChem file for each CID was obtained using the Python module PubChemPy V1.0.4, and the corresponding isomeric SMILES was used to calculate their molecular descriptors. Three of the validation compounds did not return exact PubChem matches and were excluded from filtering analysis. The remaining candidate sets were then used for filtering evaluation and receiver operating characteristic curve (ROC) analysis.

## Results and Discussion

### CCSP Performance Evaluation and Comparison Against Other Algorithms

CCSP 2.0 provides a ML approach to predicting CCS values easily and accurately in a flexible and user-modifiable environment as illustrated in Figure 1. CCSP uses the Mordred package to calculate a rich set of molecular descriptors (MD) that is more comprehensive than the MD sets obtainable using either Rdkit or Cdkit. The number of MD produced by Mordred, Rdkit and Cdkit is 1613, 115 and 258, respectively. Some prior ML CCS prediction approaches have relied on commercial packages such as Dragon 7 for MD calculations, but these packages can be costly and not supported by the vendors following release. Another key aspect of CCSP 2.0 is the reliance on a linear kernel SVR model rather than a radial basis one, making the results less prone to overfitting, more easily interpretable, and faster to calculate. The use of a faster linear kernel approach allows the user to train SVR CCS predictive models on demand, which would otherwise be time-prohibitive with a more computationally expensive approach. The SVR models in CCSP 2.0 use L2 regularization to penalize those models with higher MD weights against simpler models with lower weights overall, with the net effect being less chances of overfitting. The user’s ability of changing training sets and re-building SVR models for their own unique applications, also results in repeating the grid search step for each new SVR model built, leading to optimized epsilon (*ϵ*) and cost I hyperparameters that can compensate for higher noise values in a given data set.^25^

**Figure 1:**
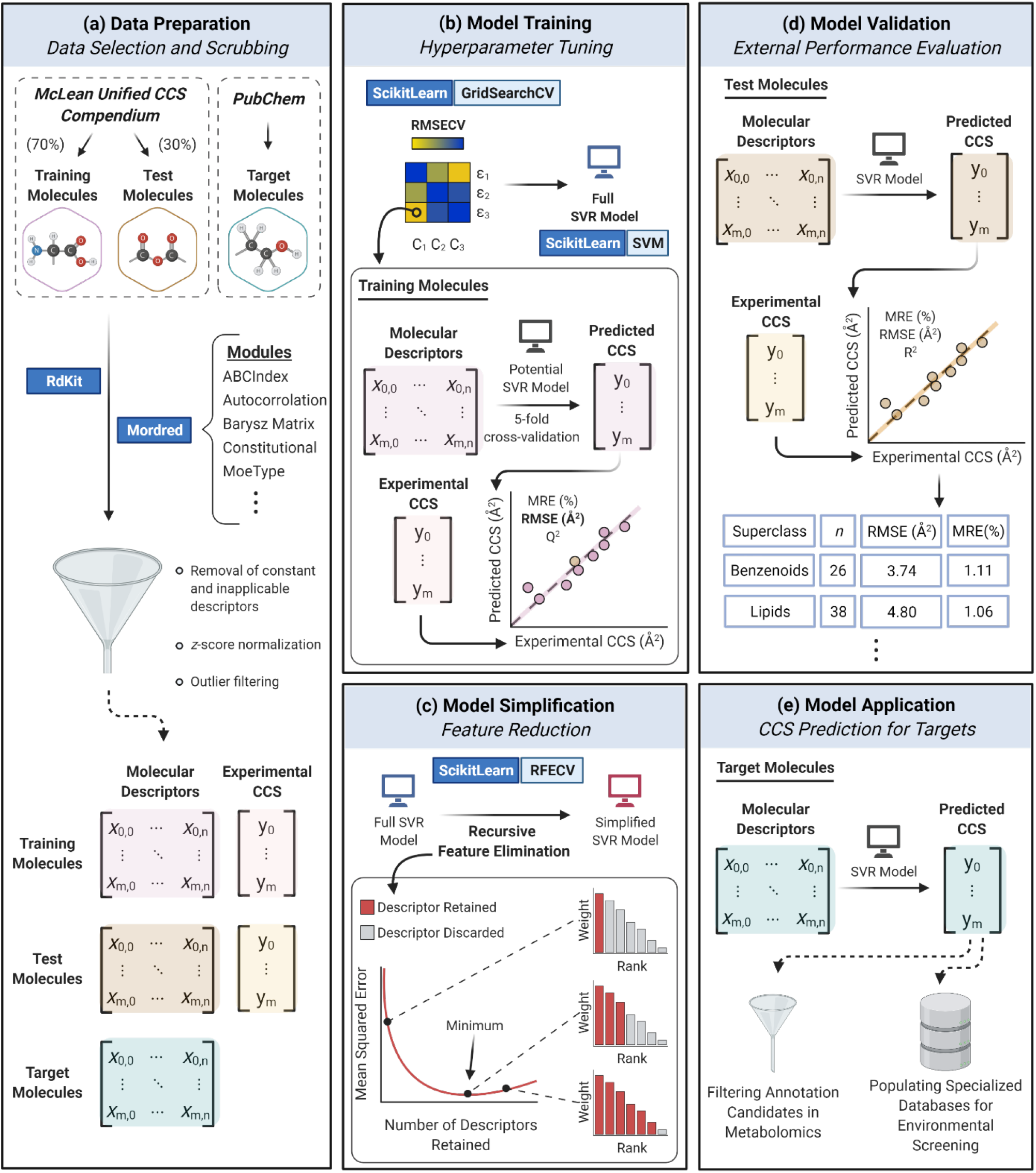
CCSP 2.0 Workflow. Graphical illustration of CCS prediction methods using user-selected training sets with a linear support vector regression (SVR) algorithm: (a) obtain reference records and prepare training and test matrices; (b) optimize model hyperparameters and perform cross-validation; (c) simplify model using recursive feature elimination with cross-validation; (d) perform model validation on withheld test set; (e) apply model to predict CCS of novel targets.

As a first step in measuring the performance of CCSP 2.0, we conducted a detailed CCS prediction accuracy study under both cross-validation and test set validation conditions. A total of 101 SVR models were created for the data sets of each adduct ion type using a different allocation of molecules for both the training (calibration) and test (validation) sets. These data sets were highly heterogeneous in terms of chemical classes, as shown in Figure S1. The root mean squared error of validation (RMSEV) was calculated for each model and models with the median RMSEV were selected for further analysis (Figure 2). Because the differential allocation of molecules to training and test sets can lead to different apparent prediction accuracies, the data split yielding the median RMSEV was selected for closer inspection to ensure fair assessment. To summarize the prediction accuracies for CCSP 2.0, global performance was evaluated using RMSEV and median relative error (MRE), while cumulative probabilities within 3 and 5% errors were used to describe the spread of prediction errors. The [M-H]^-^ model yielded a median prediction error of 1.251 % with a RMSEV of 4.998 Å^2^. Of the 170 compounds in the [M-H]^-^ validation set, 78 % of predictions fell within 3 % error, and 92 % of predictions were within 5% error. In the [M+H]^+^ SVR model, a median prediction error of 1.729 % and a RMSEV of 5.127 Å^2^ were observed. Of the 155 validation compound predictions, 70 % fell within 3 % error and 92 % were within 5 % error. Finally, the median prediction error and RMSEV for the [M+Na]^+^ validation set were 1.874 % and 6.612 Å^2^, respectively. Of the 138 [M+Na]^+^ validation predictions, 74 % fell within 3 % error and 90 % were within 5% error. Our previous version of CCSP using partial least squares regression (PLSR) models^26^ yielded MRE and RMSEV values of 1.47 % and 5.44 Å^2^ for [M-H]^-^ ions, while it produced MRE and RMSEV values of 1.84 % and 6.98 Å^2^ for [M+H]^+^ ions. Though CCSP 2.0 was evaluated using a more comprehensive CCS library than CCSP 1.0, CCSP 2.0 consistently outperformed its predecessor for similarly heterogeneous data sets. The primary advantage of CCSP 2.0 over its predecessor is its composition using one coding language (Python) and its containment to a single environment (Jupyter Notebook) for all calculations.

**Figure 2.**
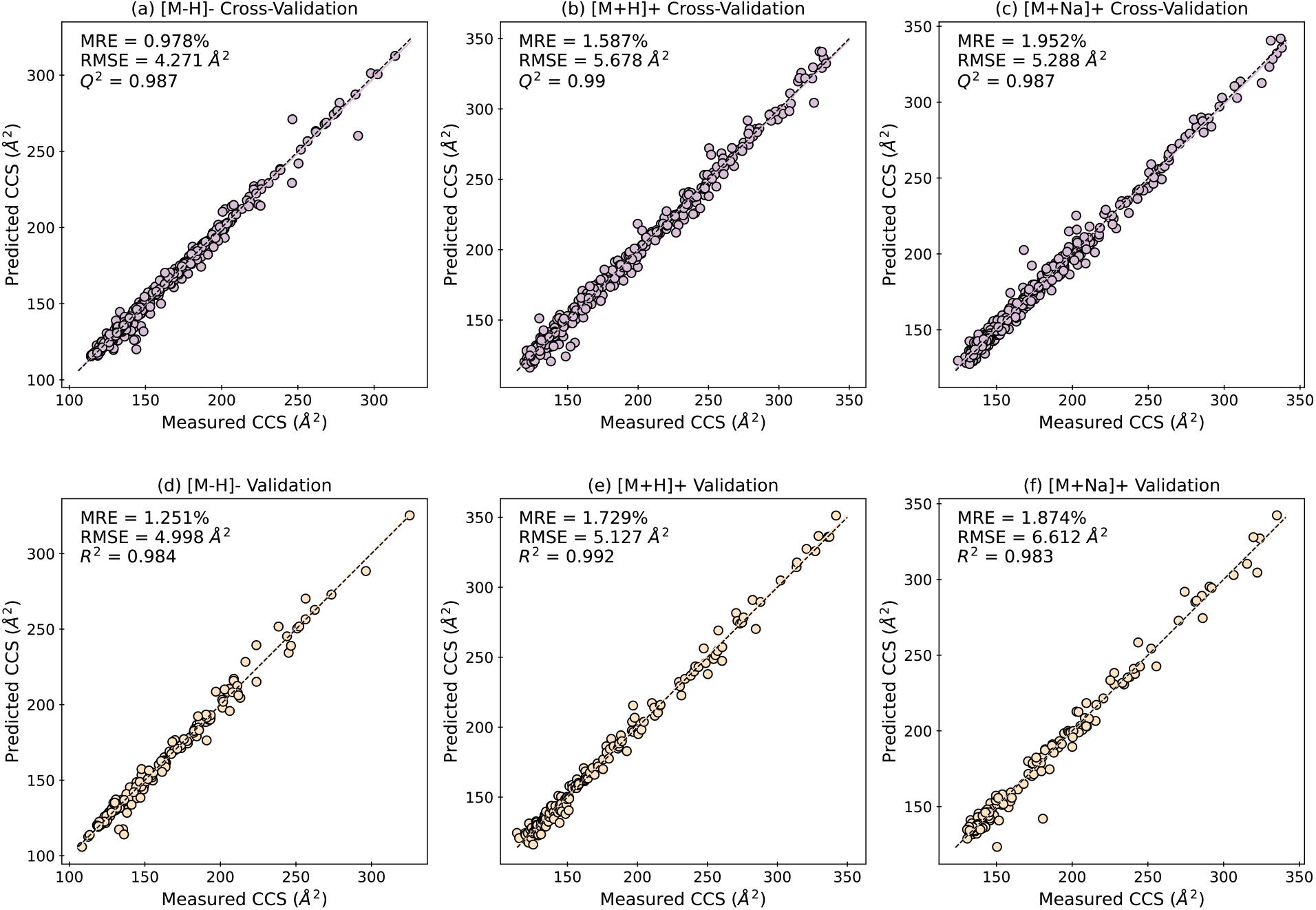
Evaluation of CCSP 2.0 prediction performance for three types of ionic species. A 5-fold cross-validation scheme was used to evaluate the performance of adduct-specific SVR models for [M-H]^-^, [M+H]^+^, and [M+Na]^+^ species (a-c). Then, independent test sets were used for the external validation of such models (d-f). The median relative percent error (“MRE”), root mean square error (“RMSE”) and coefficient of determination (“R^2^”) or goodness of prediction (“Q^2^”) are noted for each evaluation. Black dashed lines represent ideal lines of best fit, and the colored lines represent regression lines for the data set.

Although CCSP 1.0 had some of the best reported CCS prediction accuracies when compared to its contemporary approaches, its reliance on several commercial software platforms such as Dragon (Kode ChemoInformatics), Matlab (MathWorks) and the PLS Toolbox (Eigenvector) decreased its usability, as the implementation cost could be outside of what is realistic for many academic laboratories. It also required the manual download and curation of structure data files for each molecule in the training set. An appealing aspect of several current CCS prediction tools such as CCSbase and AllCCS is the availability of web servers that only require the user to input a SMILES specification to obtain a predicted CCS value. Unfortunately, the training sets are typically not user-selectable and many times the results for the individual ML models are not easily accessible for critical inspection. While Deep CCS is very flexible and allows the user to select a training set, its command-line Python interface is not the most user-friendly, particularly with trainees taking their first steps in this field. To quantitatively compare the various ML approaches in existence, we conducted a side-by-side comparison of CCSP 2.0, CCSbase, AllCCS and DeepCCS in terms of predicted CCS MRE and RMSE (Figure 3). Overall, CCSP 2.0 outperformed CCSbase, AllCCS and DeepCCS according to MRE and RMSE for the [M-H]^-^ data set for most of the 101 data splits tested. The ranking according to median MRE was CCSP 2.0 (1.251%), CCSbase (1.427%), AllCCS (1.855%), then DeepCCS (1.871%). According to median RMSE, the ranking was CCSP 2.0 (4.876 Å^2^), DeepCCS (6.068 Å^2^), CCSbase (7.359 Å^2^), then AllCCS (9.919 Å^2^). To understand the reasons behind these differences in CCS prediction accuracy, we computed the prediction accuracy for each chemical super class in each test set. The results of such analysis are presented in Figure S2, indicating that the main chemical super class where CCSP outperforms the closest competitor, CCSbase and other algorithms such as AllCCS and DeepCCS for [M-H]^-^ ions is the benzenoid superclass. The reason behind the better performance of CCSP vs. CCSbase for this chemical class seems to be the lack of shape-dependent MD as discussed below. Other chemical classes where CCSP 2.0 performed very well were the lipids and lipid-like compounds and organic acids and derivatives (Figure S2). Because these three chemical classes comprised 59% of all compounds in the data sets, CCSP had an overall better performance when compared with the other algorithms tested here.

**Figure 3.**
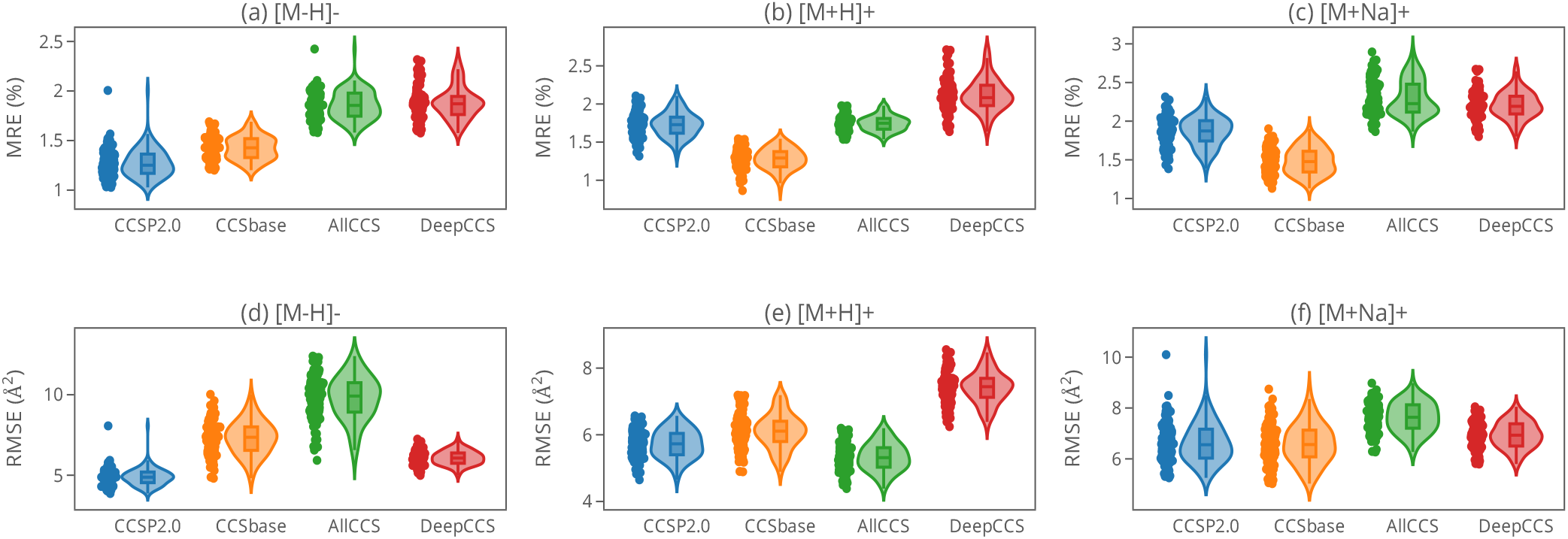
Comparison of machine learning-based CCS prediction algorithms. The performance of CCS prediction by CCSP 2.0 was compared to that of three alternative algorithms in the literature. Three data sets ([M-H]^−^, [M + H]^+^, and [M + Na]^+^) were obtained from the McLean Unified CCS Compendium and filtered based on their compatibility with all tested algorithms. The median relative percent error (MRE) and root mean square error (RMSE) were then calculated between the experimental CCS values and the CCS values predicted by each algorithm. This process was completed 101 times for each adduct ion type, each using a randomly-selected test set. Violin plots for the distribution of the MRE (a-c) and RMSE (d-e) values are provided, with the colored dots representing the results of the individual trials.

For the [M+H]^+^ data set, the ranking according to median MRE was CCSbase (1.291%), CCSP 2.0 (1.729%), AllCCS (1.748%), then DeepCCS (2.084%). According to median RMSE, the ranking was AllCCS (5.315 Å^2^), CCSP 2.0 (5.755 Å^2^), CCSbase (6.109 Å^2^), then DeepCCS (7.443 Å^2^). It was noted that while CCSbase minimized the MRE for most compound super classes investigated, the better prediction of outlier compounds by AllCCS led to a better RMSE. CCSP 2.0 balanced the two objective functions, coming in second in both rankings.

Predictions for sodiated adducts generally produced the highest prediction errors for the tools investigated, with CCSbase being the most successful. The ranking according to median MRE for the [M+Na]^+^ data set was CCSbase (1.477%), CCSP 2.0 (1.874 %), DeepCCS (2.193%), then AllCCS (2.227%). According to RMSE, the ranking was CCSP 2.0 (6.556 Å^2^), CCSbase (6.563 Å^2^), DeepCCS (6.933 Å^2^), then AllCCS (7.637 Å^2^). Notably, both CCSP 2.0 and CCSbase perform better on these species than AllCCS, the most popular CCS prediction tool.

### Effect of Molecular Descriptor Set Composition on CCS Prediction Accuracy

The SVR models used for the predictions described above were based on a selection of MD through RFE. For the [M-H]^-^, [M+H]^+^ and [M+Na]^+^ ion types, the model RMSECV exponentially decreased until approximately 50 or more descriptors were included in the calculations (Figure S3). The average number of descriptors RFE-selected under cross-validation conditions for these ion types were 370, 303 and 414, respectively. These MD were selected from the larger set of 1613 MD calculated by the Mordred package. Although the optimum SVR models used 300-400 descriptors on average, it was noted that many of these descriptors were assigned low feature weights due to the regularization scheme selected. These results explain the robust nature of the models produced by CCSP.

The most prominent descriptor classes used in SVR models (Figures S4-S6) mapped the relative position of atoms and bonds in space, particularly the Moreau-Broto autocorrelation of topological structure (ATS) descriptors, weighted Barysz Matrix descriptors, and molecular symmetry descriptors. ATS descriptors are based on graph theory and describe the distribution of atomic properties along the topological structure of a molecule.^27^ Barysz Matrix descriptors summarily describe a topological, vertex-distance matrix representation of heterosystems.^28^ Both ATS and Barysz Matrix descriptor classes can be weighted on the basis of multiple atomic properties, including electronegativity, mass, polarizability, and ionizability. The most conserved Barysz descriptors aggregated Barysz matrix eigenvectors (VE_1_ and VE_3_ series) or Randic-like^29^ eigenvectors (VR_2_ and VR_3_ series), while the most retained ATS descriptors described the distribution of atomic properties for atoms close in space (ATS_0_, ATS_1_, ATS_2_ and ATS_3_ series). The Kappa Shape Index (Kier)^30^ and Total Information Content (TIC)^31^ descriptors, which encode molecular symmetry, were also highly conserved in SVR models and enabled CCSP 2.0 to consider cis-trans isomerism in CCS predictions. Descriptors that predict the bulk volume (*e.g*. Van der Waals volume (V_ABC_) and McGowan Volume (V_Mc_)) and those that describe bulk polarizability (*e.g*. Crippen-Wildman molar refractivity (SMR) and atomic polarizability (apol)) were also often conserved. The last highly retained set of descriptors belonged to the Mordred constitutional module, which included the sum of atomic properties relative to the ^12^C properties. Notably, these descriptors were previously found to be critical to our previous CCS prediction approach, CCSP 1.0, which used a different ML approach and employed licensed software to calculate >5000 descriptors.^26^

To examine if the set of most conserved Mordred descriptors would be sufficient to produce robust and highly accurate SVR models to predict CCS values, we investigated three separate variable selection schemes (Figure S7). A set of “consensus” descriptors was created by selecting those MD that were conserved in at least 95% of SVR models tested for each ion-type. Results indicated that the consensus MD set did not provide significantly better CCS accuracies when compared against the set containing all descriptors, or the set of RFE-selected descriptors. The list of consensus descriptors tested in these models is provided as a supplemental spreadsheet (Supplemental Table 6). Though smaller MD sets may make for simpler models if desired, these results show that they do not afford better prediction accuracy. In conjunction with the similar errors between the cross-validation and validation rounds, these results show that the L2 regularization scheme was able to avoid overfitting, which may be a concern when using a large number of molecular descriptors.

### CCSP Applications to Metabolite Annotation

As a first test of the capabilities of CCSP 2.0 to predict CCS values for the purposes of filtering false positive annotations of metabolites and other small molecules, we chose an ion with a relatively small *m/z* value (isoorotic acid, [M-H]^−^ = 155.0093). The PubChem database was then searched for all candidate [M-H]^−^ ions that matched the *m/z* within a 10 ppm tolerance. The choice of database was motivated to simulate the worse-case scenario where the general chemical class of the analyte being annotated is completely unknown. We chose PubChem *vs*. more focused databases such as HMDB^32^, so as to test the CCS filtering approach in the most adverse conditions.

We chose a mass error of 10 ppm for database searches to a) compare against previous work with DeepCCS and similar algorithms and b) accommodate the realistic everyday mass accuracy of typical mass spectrometers coupled with IM, such as time-of-flight (ToF) MS. Figure 4 summarizes the results obtained for this case study. The PubChem query for *m/z* 155.0093 returned a total of 221 single-compound candidates within a 10 ppm mass window. CCS values for all these 221 candidates were predicted with CCSP (Figure S8), and only those candidates (*n* = 32) within a ±2 % window were retained, demonstrating the power of CCS filtering to remove false positive matches (86 %) to an unknown compound being annotated. As expected, when the mass search threshold was decreased from 10 ppm to lower settings, the number of compounds matching the unknown also decreased, emphasizing the importance of simultaneous good accuracy in both the *m/z* and CCS dimensions. As the mass tolerance was made smaller, a lower percentage of candidates was filtered by CCS. The experimentally-measured CCS value for isoorotic acid was 120.22 Å^2^, and the CCSP-predicted value was 119.3 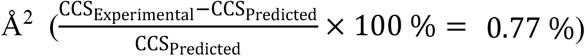 In terms of CCS confidence threshold to keep the correct annotation candidates, we chose a 2% limit, with literature values ranging 1-4%. The MRE for the SVR model used for prediction of CCS values was 1.25 % (Figure 2), so the ±2% threshold is somewhat conservative. Interestingly, CCS measurements for the two isobaric species orotic and isoorotic acid, with predicted CCS values of 119.3 Å^2^ and 121.7 Å^2^ 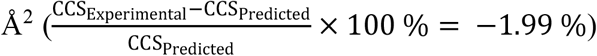 may not be adequately distinguished with a 2% threshold, depending on rounding, but would be distinguished well with a less conservative threshold of 1.5%, for example.

**Figure 4:**
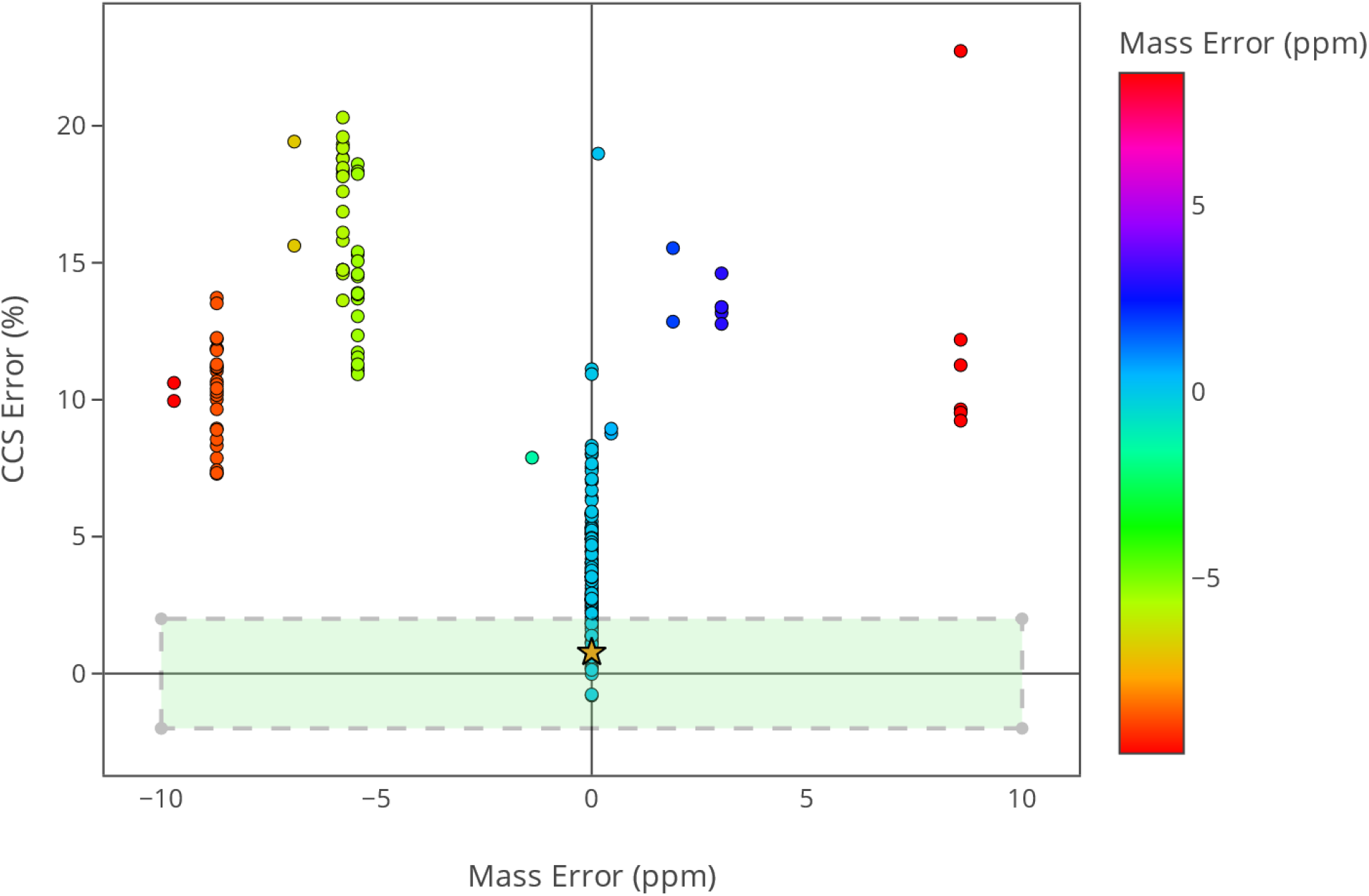
Effect of mass and CCS accuracy on the number of filtered metabolite annotation candidates. The use of CCS in ion mobility-mass spectrometry studies for the purpose of filtering candidate molecules was evaluated by treating isoorotic acid (C_5_H_4_N_2_O_4_) as an unknown species. A PubChem query returned 221 candidate molecules with monoisotopic masses within 10 ppm of the theoretical monoisotopic mass of isoorotic acid. The CCS value of each of the 221 queried molecules was predicted using a SVR model developed with CCSP 2.0. This model was trained with 70% of the [M-H]^-^ ions in the McLean Compendium, chosen randomly (Supplemental Table 4). The green shaded box depicts the joint ±10 ppm and ±2 % CCS interval surrounding the experimental CCS of isoorotic acid (119.3 Å^2^) in the [M-H]^-^ form.

To evaluate the impact of the CCS filtering threshold (Δ_CCS_) on the percentage of filtered compounds for various mass accuracies, the workflow used in the isoorotic acid case study was applied to 162 deprotonated test compounds. At a mass tolerance of 10 ppm, PubChem searches for each test compound returned between 15 and 18,399 candidates for a combined total of 598,208 candidates. Figure 5 illustrates the decrease in the percent of filtered annotation(s) as the CCS filtering threshold is increased from 0-15%. Conversely, as the Δ_CCS_ is made smaller, more incorrect annotation candidates are filtered, but the risk of over-filtering the right annotation also increases. At Δ_CCS_ 3%, 76.5% of the true annotations for the test compounds were retained, and a median of 47.3% of the candidates were filtered for lists generated with a 10 ppm mass window. The median percentage of candidates reduced drops to 24.9% when the candidates are restricted to only those with the same chemical formula as the true species. This analysis reveals that the choice of Δ_CCS_ threshold for filtering purposes depends on several factors and that there is a tradeoff in the number of correct annotations retained and false positive reduction. For the data in Figure 5, no single CCS filtering threshold appears effective enough to retain 100% of the correct annotations for the diverse chemical data set involved in this case example.

**Figure 5.**
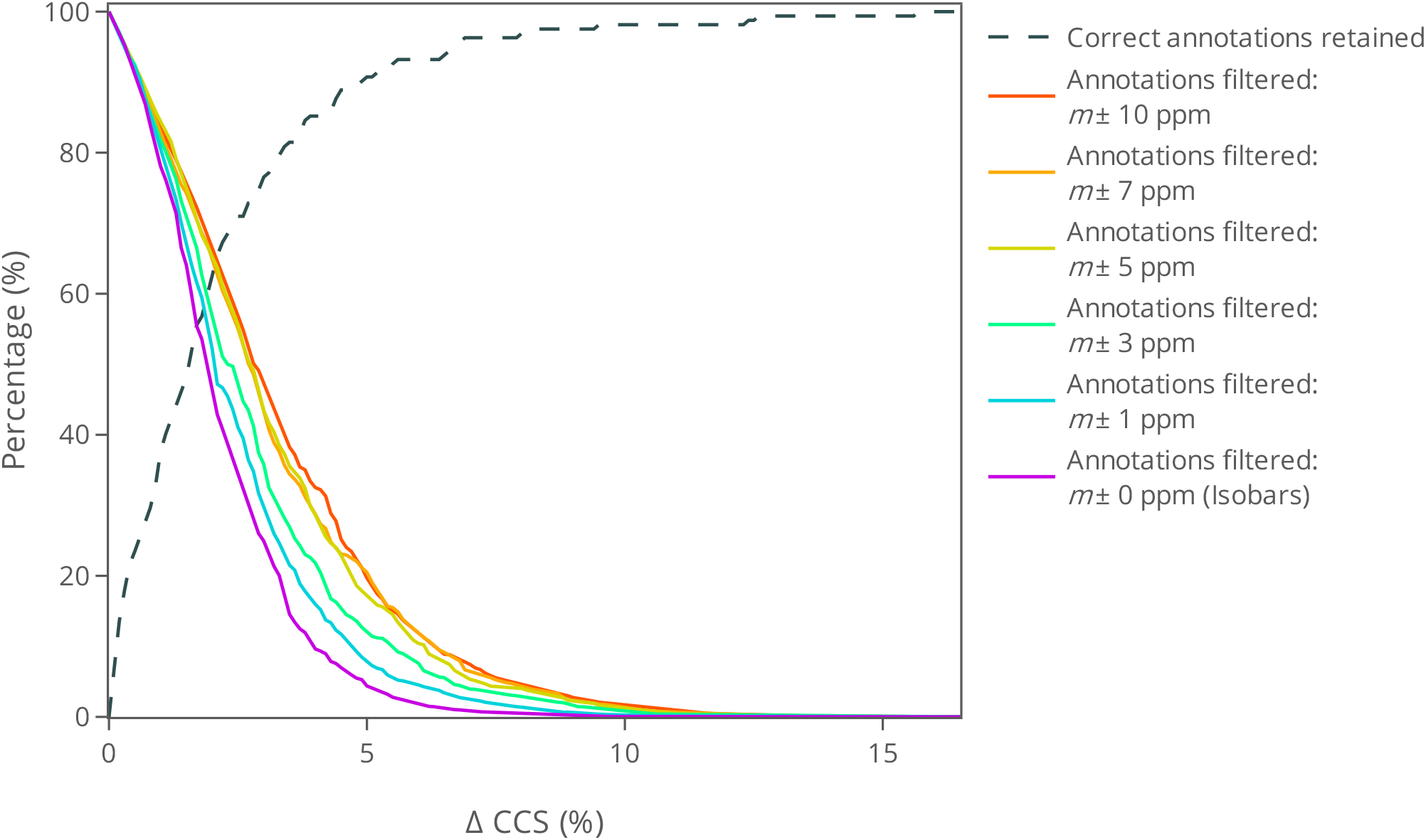
Effect of CCS accuracy on the number of filtered metabolite annotation candidates for a test set consisting of 162 [M-H]^-^ species. A test set of *n* = 162 CCS values for [M-H]^−^ions was used to evaluate the filtering efficacy of CCS (Supplemental Table 4). Each compound in the test set was treated as an unknown, with only its monoisotopic mass and experimental CCS value being utilized. For each validation compound, all entries in PubChem with monoisotopic masses within 10 ppm of its monoisotopic mass (*m*) were retrieved. The CCS value of each PubChem candidate was predicted with CCSP 2.0, using *n* = 370 training molecules and the whole set of Mordred descriptors. The grey dashed line represents the percentage of the correct compound structures in the test set whose predicted CCS value fell within the set %CCS tolerance. The remaining traces represent the median percentage of candidates that were filtered from each candidate list at a specified % CCS at a given mass tolerance. The experiment was performed for candidate lists generated with 10 ppm (red), 7 ppm (orange), 5 ppm (gold), 3 ppm (green), 1 ppm (blue), and 0 ppm (purple) mass thresholds.

A more comprehensive approach to examine annotation success by combining CCS and accurate mass values is using receiver operating characteristic (ROC) curves.^33^ Rather than rely on individual Δ_CCS_ thresholds, the ROC curve plots true positive discovery rates *vs*. false positive discovery rates at multiple Δ_CCS_ thresholds. The area under the ROC curve (AUC) is a metric for the algorithm’s filtering efficacy, with an AUC of 1 representing a method capable of rejecting all incorrect annotations while retaining all correct annotations. Figure S9 reveals that decreasing the mass window used to generate the annotation candidate lists generally reduces the AUC, with CCS filtering failing to meaningfully differentiate correct annotations from incorrect annotations with the same chemical formula. The AUC scores for candidate lists based on 10 ppm, 7 ppm, 5 ppm, 3 ppm, 1 ppm, and 0 ppm mass windows were 0.657, 0.633, 0.622, 0.616, 0.593, and 0.589, respectively. These results suggest that IM-MS systems likely afford little CCS filtering capacity if used in conjunction with CCS prediction algorithms that are built on chemically diverse training sets. These chemically diverse training sets produce CCS predictions that may not be accurate enough to filter sufficient candidates. However, if the CCS prediction accuracy is improved using a training set that is structurally closer to the analyte, then CCS filtering produces more satisfactory results, as seen in prior predictions for structurally-similar xenobiotics^34^. Notably, the effectiveness of CCS filtering also depends on the mass accuracy afforded by the back-end mass spectrometer. For highly accurate mass spectrometers, lower CCS prediction errors and more accurate CCS experimental measurements are required to make the CCS filtering approach more valuable.

To illustrate the utility of CCS prediction using a structurally similar training set, we investigated deprotonated species of the lipid and lipid-like superclass. This superclass includes molecular classes such as fatty acyls, prenol lipids, glycerolipids, glycerophospholipids, sphingolipids, steroids and steroid derivatives. Figures S10 and 6 depict a case study involving a lipid species, leukotriene E4. In this case, CCSP was trained only with molecules in the lipid superclass, leading to more accurate CCS predictions than if the model was trained with a more diverse range of molecules. Out of the 4,980 molecules queried from PubChem within a 10 ppm window of the analyte, only 806 remained using a ±2 % CCS filtering interval. This candidate space could be further reduced by improving mass accuracy, as discussed below. Figure 6 demonstrates that the presence of halogen atoms such as chlorine or fluorine reduce the predicted CCS of the annotation candidates, while the presence of silicon increases the predicted CCS values. These atom types frequently appear in lipid candidate lists generated by mass and may contribute to the improved filtering of annotation candidates in lipid case studies. Though chlorinated, fluorinated and silconated species made up 5%, 21%, and 2% of the annotation candidates in this case study, respectively, the impact of specialized training set extended even to isobaric lipid candidates. Approximately 57 % of the isomers of leukotriene E4 could be filtered with a ±2 % CCS filtering interval, demonstrating that CCS prediction could improve confidence in the isobaric annotation of the lipid mediator in addition to rapidly rejecting false positive matches to halogenated and siliconated species.

**Figure 6.**
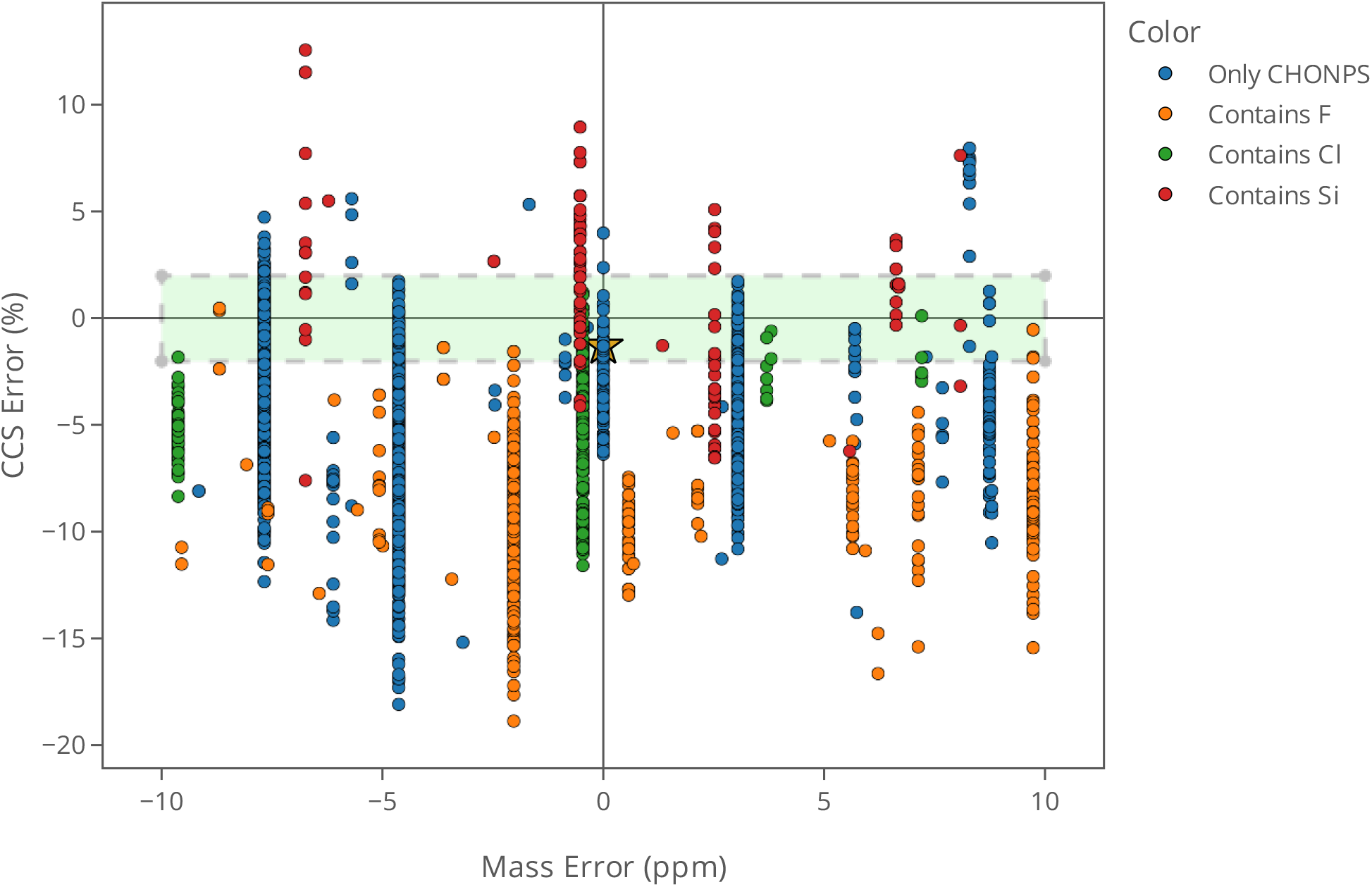
Effect of mass and CCS accuracy on the number of filtered metabolite annotation candidates for the case of a lipid molecule. The use of CCS in ion mobility-mass spectrometry studies for the purpose of filtering candidate molecules was evaluated by treating one randomly-selected compound in the validation set, leukotriene E4, as an unknown species. The only information utilized for this compound was the theoretical monoisotopic *m/z* of the [M-H]^-^ ion (C_23_H_38_NO_5_S) and the experimental CCS value (213.2 Å^2^) listed in the McLean Unified CCS Compendium. A PubChem query returned 4980 candidate molecules with monoisotopic masses within 10 ppm of the theoretical monoisotopic mass of leukotriene E4. The CCS value of each of the 4980 queried molecules was predicted using a SVR model developed with CCSP 2.0. This model was trained with 93 [M-H]^-^ ions in the lipid superclass of the McLean Compendium (Supplemental Table 5). The green shaded box depicts the joint ±10 ppm and ±2 % CCS interval. The various PubChem candidates are colored based on the presence of various types of atoms other than CHNOPS in their structures.

Figure 7 shows the effect of CCS filtering threshold on the percentage of retained correct annotations and % of candidates filtered when CCSP SVR models were trained with 93 [M-H]^-^ ions in the lipid superclass of the McLean CCS Compendium and tested on 33 compounds in that class. At a mass tolerance of 10 ppm, PubChem searches for each test compound returned between 43 and 13,984 candidates for a combined total of 136,940 candidates. At a Δ_CCS_ threshold of 2.8%, a total of 100% of the correct annotations were retained, again emphasizing the importance of using a structure-matched training set in ML CCS prediction. At this Δ_CCS_ threshold, a median of 36.1% of candidates generated with a mass tolerance of 10 ppm could be filtered without accidently filtering any correct annotations. At the same Δ_CCS_ threshold, 19.3% of isomeric species (0 ppm error) could be filtered. These results indicate that CCS can be used as a preliminary filter to rapidly reject a large number of potential candidates and that even truly isobaric species can benefit from CCS prediction.

**Figure 7.**
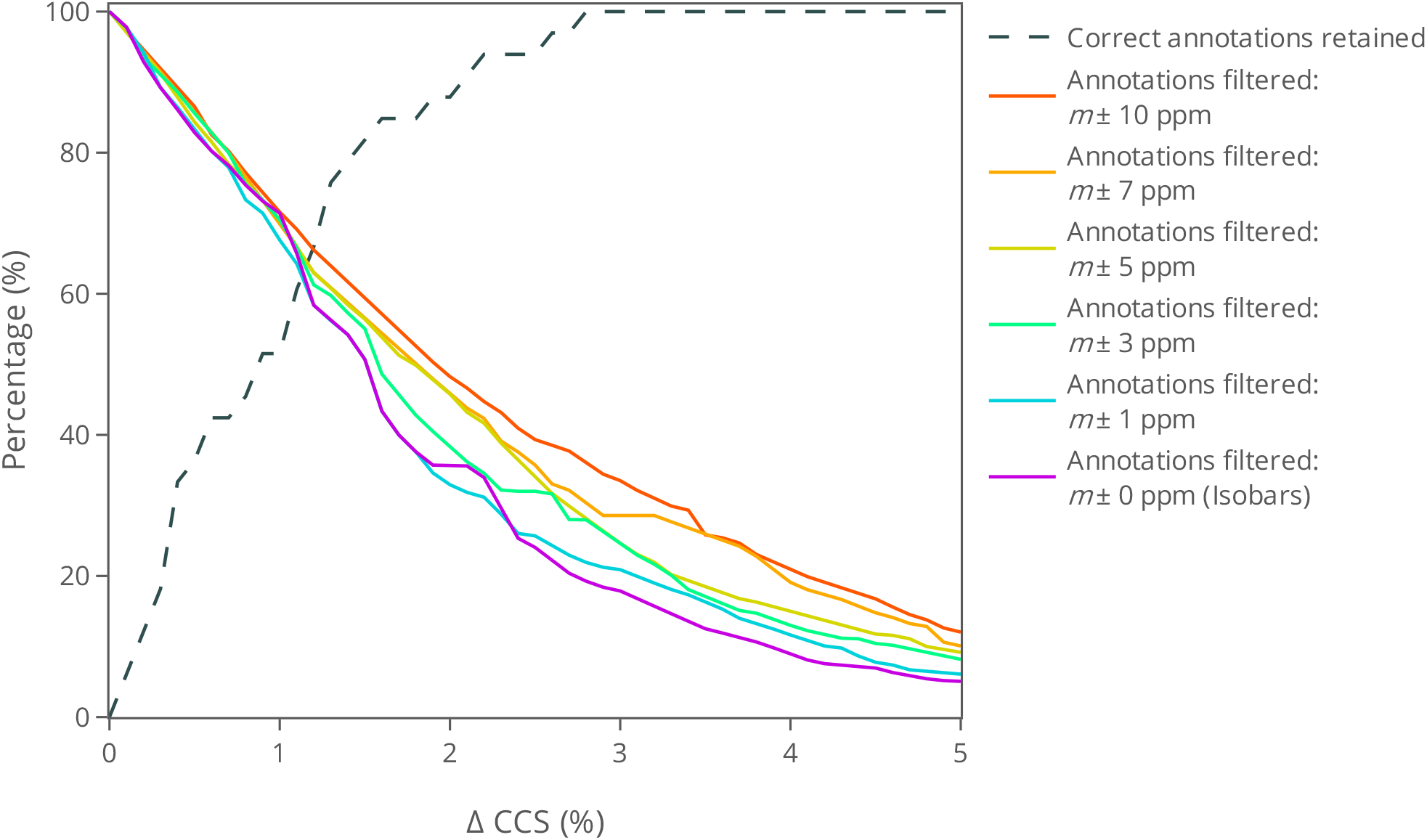
Effect of CCS accuracy on the number of filtered lipid annotation candidates for a test set consisting of 33 [M-H]^-^ lipid species. A set of *n* = 129 CCS values for [M-H]^−^ ions was obtained from the lipid superclass of the McLean Unified CCS Compendium (Supplemental Table 5). The data set was randomly split with 70% being allocated to the training of SVR models. The remaining 30% of the data was retained for evaluation of the number of lipid candidates filtered through CCS matching. Each validation compound in the test set was treated as an unknown lipid, with only its monoisotopic mass and experimental CCS value being utilized. For each validation compound, all entries in PubChem with monoisotopic masses within 10 ppm of its monoisotopic mass (*m*) were retrieved. The CCS value of each PubChem candidate was predicted using CCSP 2.0 with the whole set of molecular descriptors. The grey dashed line represents the percentage of the correct compound structures in the test set whose predicted CCS value fell within the set %CCS tolerance. The remaining traces represent the median percentage of candidates that were filtered from each candidate list at a specified % CCS at a given mass tolerance. The experiment was performed for candidate lists generated with 10 ppm (red), 7 ppm (orange), 5 ppm (gold), 3 ppm (green), 1 ppm (blue), and 0 ppm (purple) mass thresholds.

An ROC analysis for the case of lipid analytes is shown in Figure S11. AUC scores for candidate lists based on 10, 7, 5, 3, 1, and 0 (isobaric) ppm mass windows were 0.776, 0.754, 0.748, 0.768, 0.756, and 0.756, respectively. Notably, this AUC analysis shows that CCS filtering for the lipid class is similarly effective for candidate lists generated with differing mass windows, making the class-matched approach more robust towards deviations in accurate mass measurements. These results contrast the sharp decline in filtering effectiveness for narrower mass windows in the chemically diverse data set and suggest that CCS may make an adequate classifier to differentiate true positives from false positives in class-matched metabolite annotation.

## Conclusions

In this manuscript we present CCSP 2.0, a new Python algorithm in the Jupiter environment that enables CCS predictions that are better aligned with FAIR principles. Extensive testing of the CCS prediction accuracy was conducted under both cross-validation and test set validation conditions. The CCS prediction median relative errors were 1.25, 1.73 and 1.87% for [M-H]^-^, [M+H]^+^ and [M+Na]^+^ adducts using linear support vector regression models. The performance of the new algorithm was compared against three popular machine learning CCS prediction methods reported in the literature using 101 randomly-selected test sets that were kept consistent across all algorithms. Results showed that in most cases, CCSP 2.0 performed better than or equally well as other approaches, with the advantages that the training set can be easily customizable for the specific application in mind and that a larger set of molecular descriptors is available to explain more subtle structural differences in the tested analytes. Despite the outlined advantages, CCSP 2.0 still suffers from some drawbacks. These include the requirement for familiarity with a programming language such as Python and the need to define the specific training set to be used prior to predicting CCS values. Current training sets allow the prediction of the most common adduct ion types including [M-H]^-^, [M+H]^+^, [M+Na]^+^ and [M+K]^+^ but not dimers such as [2M+H]^+^ or [2M-H]^-^, or other less common adducts or in-source fragments found in ESI-MS experiments.

Our filtering studies suggest that carefully class-matched models may improve the confidence of a structural annotation, but that currently available tools afford minimal orthogonal information for diverse data sets with highly accurate mass measurements. As IM technology and calibration methods improve, more accurate CCS measurements are expected to become available and will allow the production of even more accurate predictions. Future work will include an exploration of CCS predictions across multiple IM-MS instrument types, such as drift tube and travelling wave IM platforms, and an in-depth investigation of the effect of experimental sources of variance on the choice of CCS threshold chosen for filtering annotation candidates. Additional future work will include the use of multi-adduct predictions to evaluate CCS filtering and the CCS prediction of fragment ions in tandem mass spectrometry.

## Acknowledgements

We acknowledge support by NIH 1U2CES030167-01 and 1R01CA218664-01. Elements of Figure 1 were created using tools from BioRender.com.

