## Supplemental Information for "CCS Predictor 2.0: An Open-Source Jupyter Notebook Tool for Filtering Out False Positives in Metabolomics"

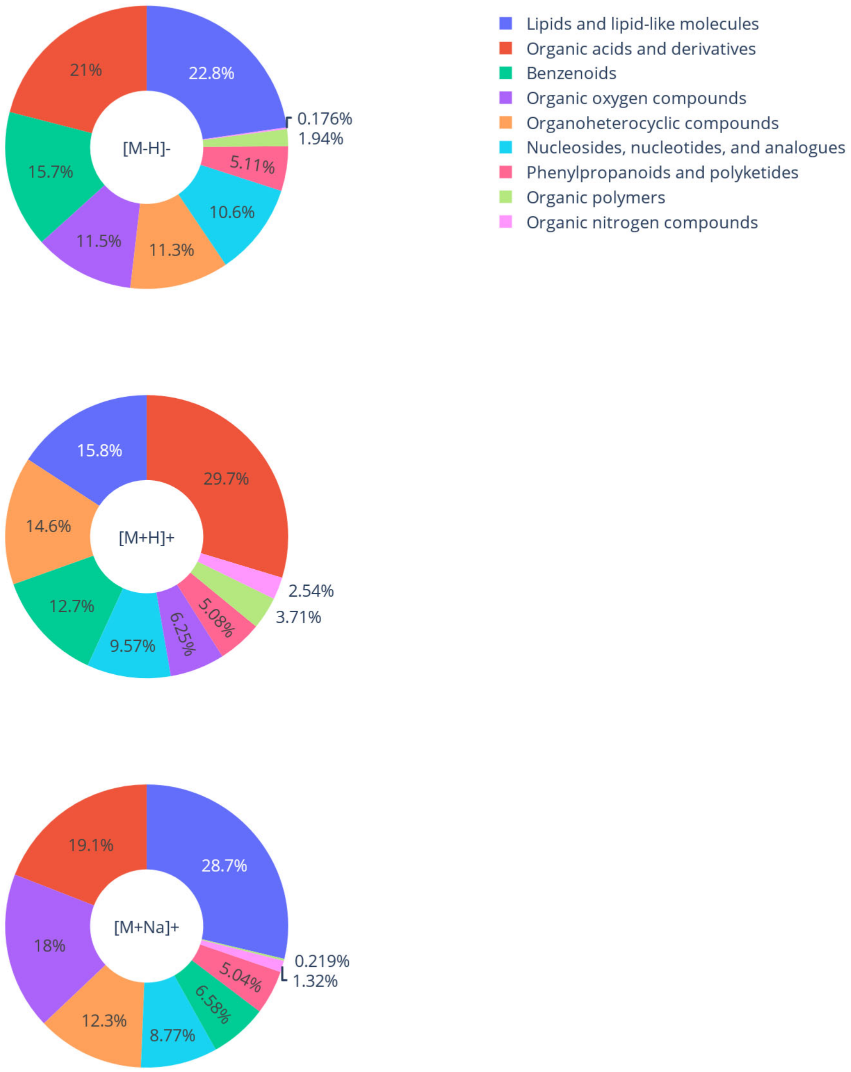


**Figure S1**: **McLean Unified CCS Compendium super class count by adduct ion-type.** CCSP 2.0 was used to predict CCS values of three chemically-heterogeneous data sets obtained from the McLean Unified CCS Compendium. These data sets were the singly deprotonated ions (top panel, $[\text{M-H}]^-$, $n = 567$), the singly protonated ions (middle panel, $[\text{M+H}]^+$, $n = 518$), and the singly sodiated ions (bottom panel, $[\text{M+Na}]^+$, $n = 461$). Pie charts represent the super class composition of each set as determined by ClassyFire[1].

.

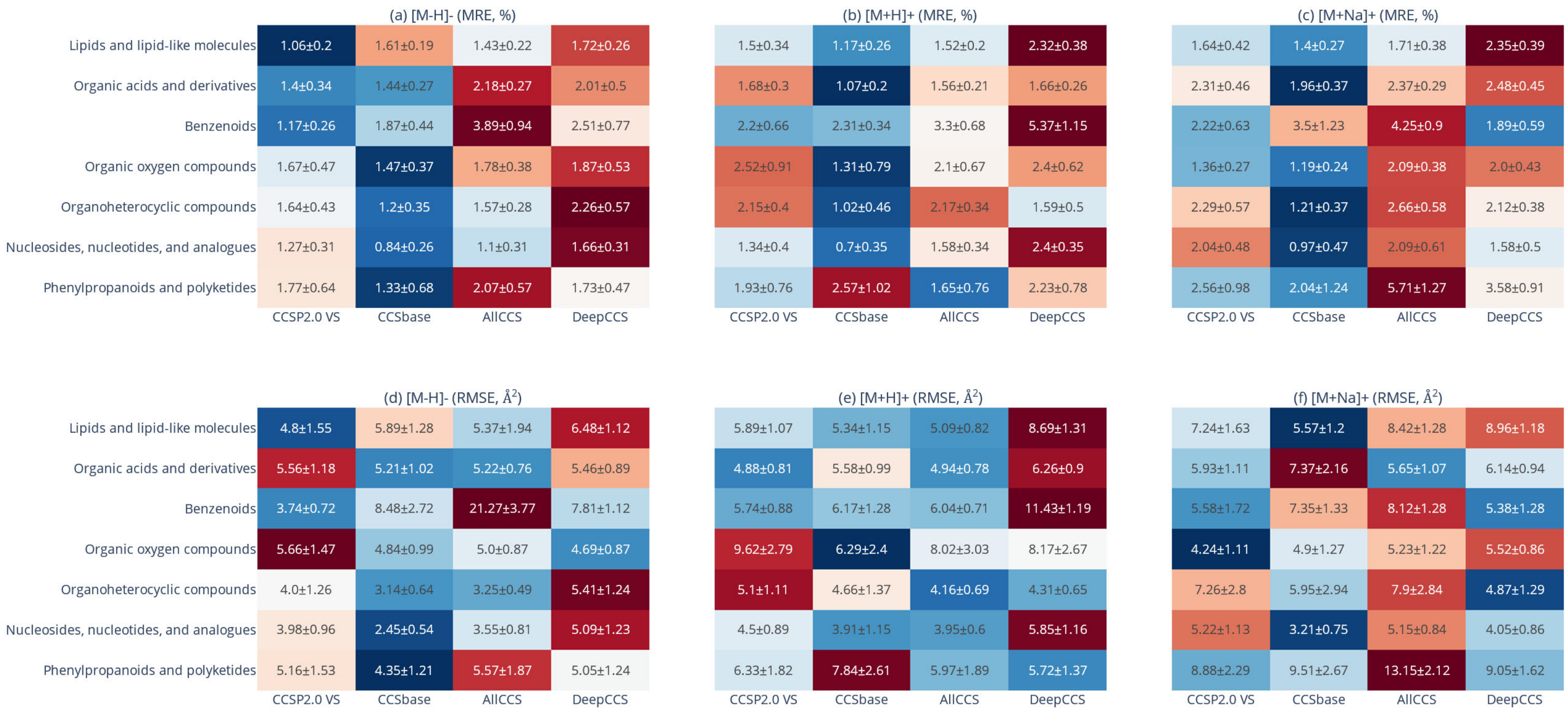


**Figure S2**: **Heatmap comparison of the performance (MRE & RMSE) of CCS prediction algorithms by adduct ion type and chemical super class.** The performance (MRE & RMSE) of CCS prediction by CCSP 2.0 was compared to that of three alternative algorithms in the literature. Three data sets were obtained from the McLean Unified CCS Compendium and filtered based on their compatibility with all tested algorithms. The $[M\text{-}H]^-$ ion set contained $n = 567$ compounds, the $[M+H]^+$ ion set contained $n = 518$ compounds, and the $[M+Na]^+$ ion set contained $n = 461$ compounds. In a single run, the CCS values of 30% of a given data set was predicted using CCSP 2.0, CCSbase, AllCCS, and DeepCCS. The median relative percent error (MRE) and root mean square error (RMSE) were then calculated between the experimental CCS values and the CCS values predicted by each algorithm. This process was completed 101 times for each ion type, with each trial using a randomly selected test set. The algorithms were parsed into the seven super classes of molecules that composed more than 5% of the data set. Heat maps display the median relative errors (a-c) and root mean square errors (d-e); the labels of each pane show the mean ± standard deviation of the 101 trials, while the colors are normalized as *Z*-score according to row.

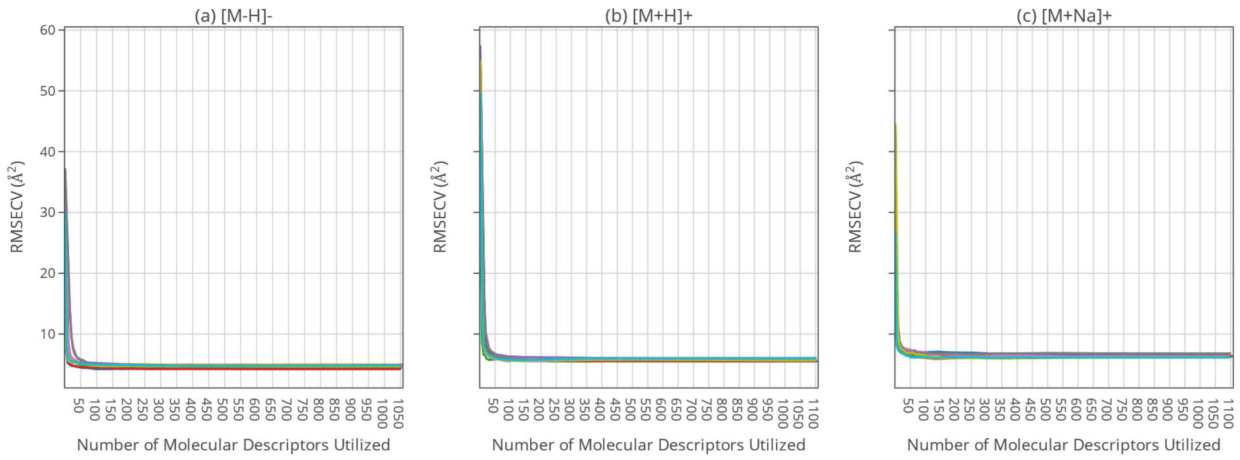


**Figure S3. Representative recursive feature elimination curves for CCSP 2.0 model calibration.** Support vector regression models were built to predict the collision cross sections of three ion-types using 2-dimensional molecular descriptors (MD) calculated by the Python package Mordred. A total of 101 models were constructed for each ion-type, where each iteration was trained using a different subset (70%) of the corresponding McLean Unified CCS Compendium entries. To reduce model complexity, each iteration employed recursive feature elimination to select the optimum number of MD that minimized the root mean square error of cross validation (RMSECV). The first ten feature selection curves are presented for the (a) $[M\text{-}H]^-$, (b) $[M+H]^+$, and (c) $[M+Na]^+$ ion-types.

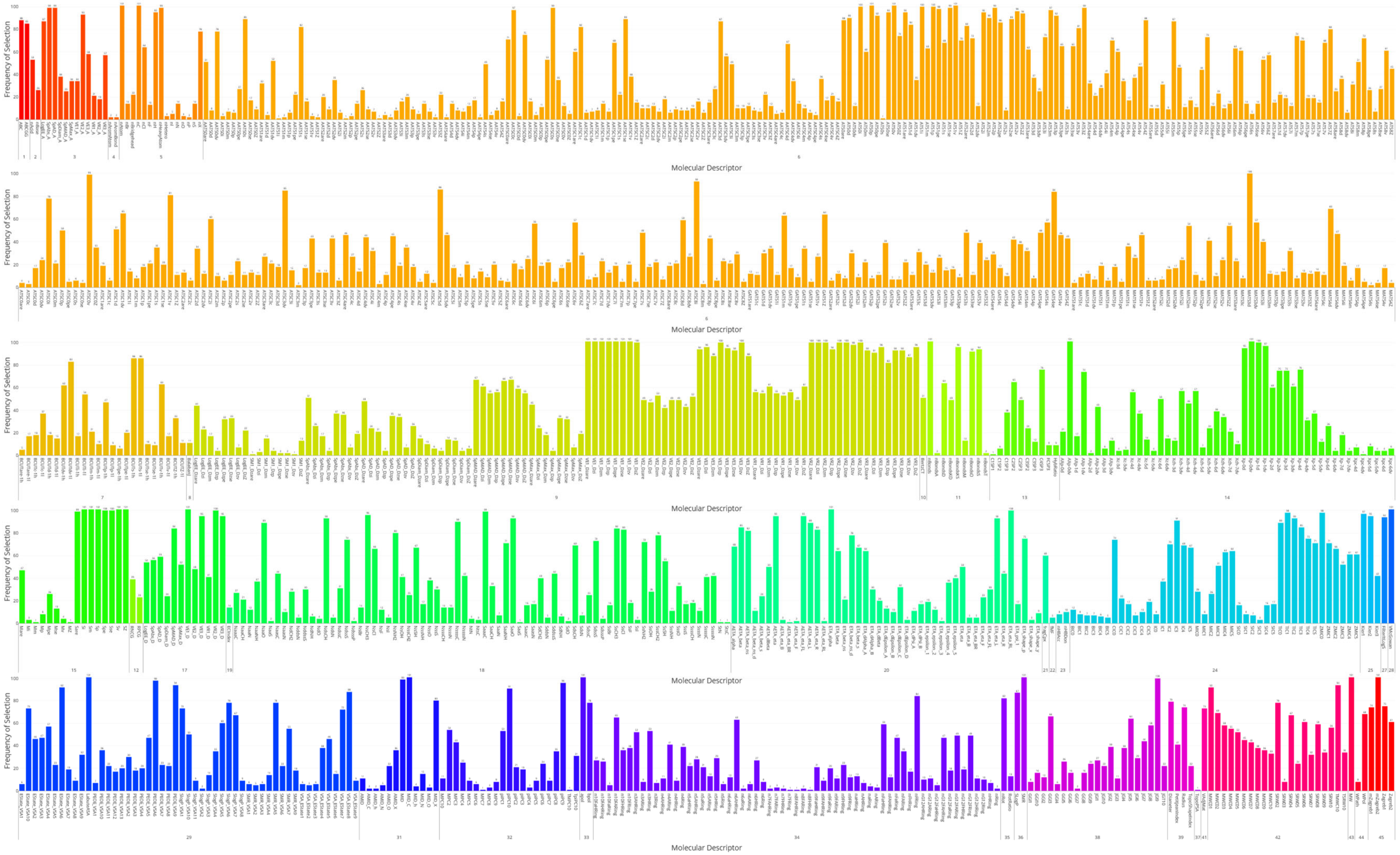


**Figure S4: Mordred molecular descriptor selection frequency for the prediction of CCS values for deprotonated molecules.** CCS predictions were performed for the $[M\text{-}H]^{-}$ ion data set obtained from the McLean Unified CCS Compendium using CCSP 2.0. The ion set contained $n = 567$ compounds, and 70% of the compounds were used to train a linear support vector regression model. The remaining 30% were held for external validation. Recursive feature elimination was used to reduce the feature space by iteratively removing the lowest weighted features until the root mean square error of cross-validation increased. The process was repeated for 101 trials each containing different data splits, selecting different feature subsets in each trial. Each vertical bar represents the number of times a descriptor was selected, omitting descriptors that were never selected. The Mordred MD modules are depicted with numbers under each set of vertical bars. A list of module names is provided in Table S7. Graph is vector-based to allow zooming.

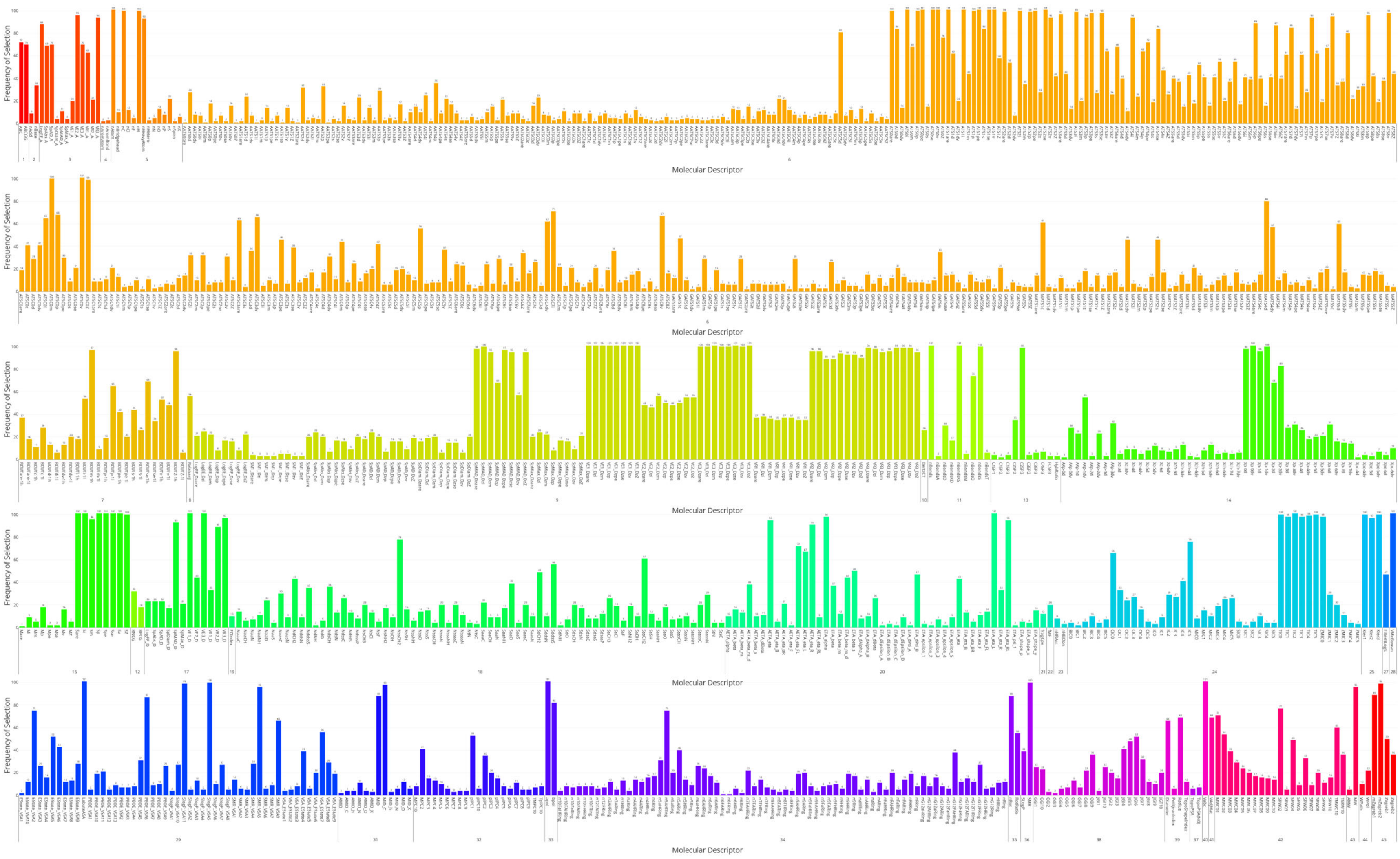


**Figure S5: Mordred molecular descriptor selection frequency for the prediction of CCS values for protonated molecules.** CCS predictions were performed for the [M-H]$^+$ ion data set obtained from the McLean Unified CCS Compendium using CCSP 2.0. The ion set contained $n =$ 518 compounds, and 70% of the compounds were used to train a linear support vector regression model. The remaining 30% were held for external validation. Recursive feature elimination was used to reduce the feature space by iteratively removing the lowest weighted features until the root mean square error of cross-validation increased. The process was repeated for 101 trials each containing different data splits, selecting different feature subsets in each trial. Each vertical bar represents the number of times a descriptor was selected, omitting descriptors that were never selected. The Mordred MD modules are depicted with numbers under each set of vertical bars. A list of module names is provided in Table S7. Graph is vector-based to allow zooming.

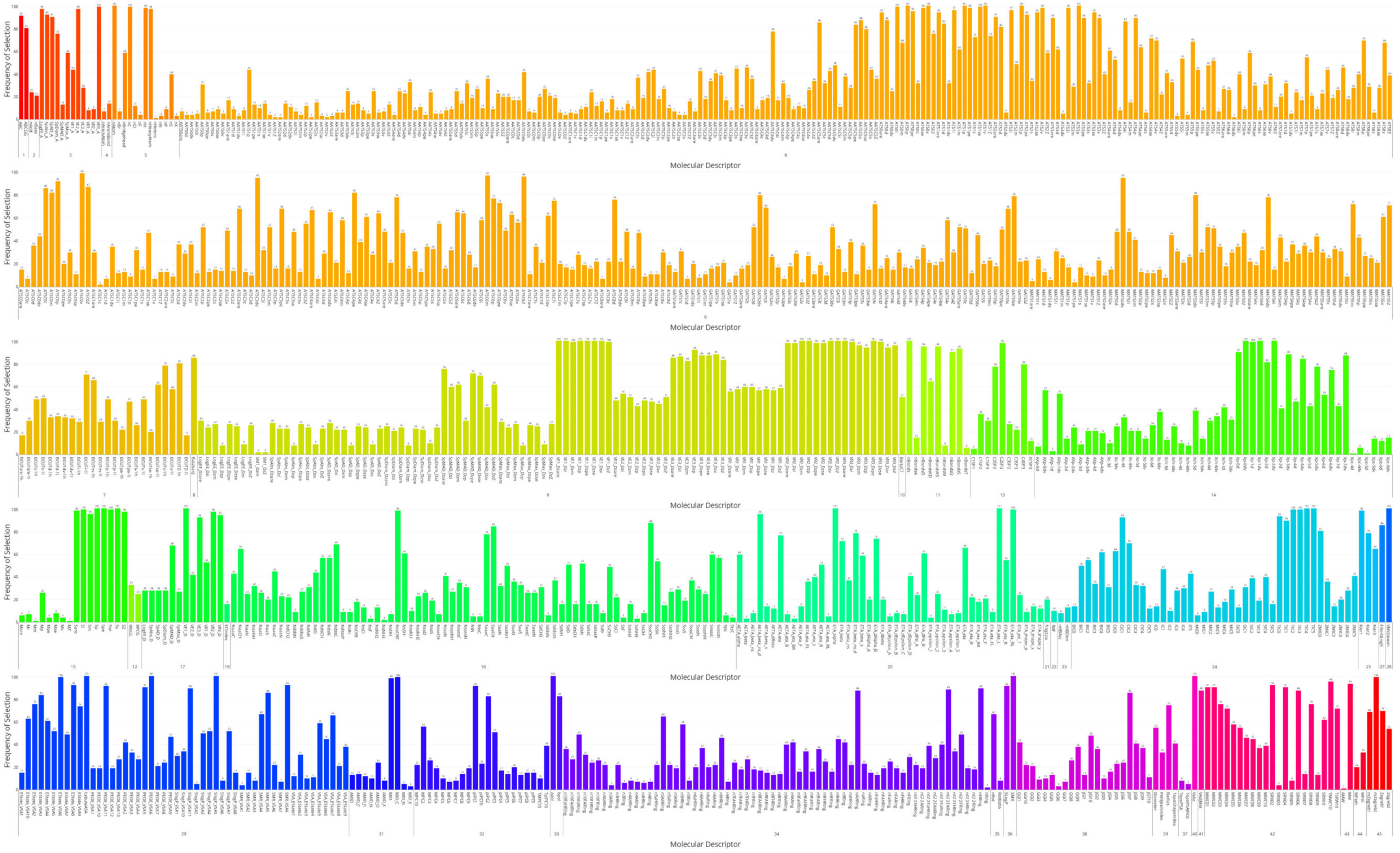


**Figure S6: Mordred molecular descriptor selection frequency for the prediction of CCS values for sodiated molecules.** CCS predictions were performed for the $[M+Na]^{+}$ ion data set obtained from the McLean Unified CCS Compendium using CCSP 2.0. The ion set contained $n = 461$ compounds, and 70% of the compounds were used to train a linear support vector regression model. The remaining 30% were held for external validation. Recursive feature elimination was used to reduce the feature space by iteratively removing the lowest weighted features until the root mean square error of cross-validation increased. The process was repeated for 101 trials each containing different data splits, selecting different feature subsets in each trial. Each vertical bar represents the number of times a descriptor was selected, omitting descriptors that were never selected. The Mordred MD modules are depicted with numbers under each set of vertical bars. A list of module names is provided in Table S7. Graph is vector-based to allow zooming.

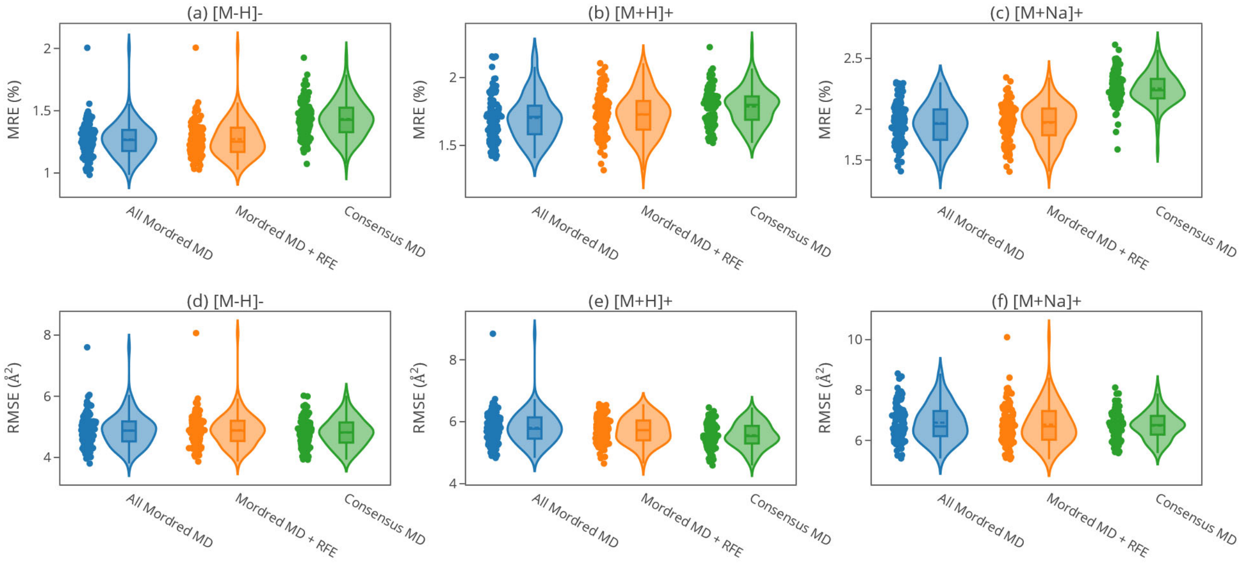


**Figure S7. Evaluation of CCSP 2.0 prediction performance using three molecular descriptor selection schemes.** CCSP 2.0 was used to predict the collision cross sections for subsets of three ion-type data sets ($[M-H]^-$, $[M+H]^+$, $[M+Na]^+$) from the McLean Unified CCS Compendium using three different molecular descriptor selection schemes. In the first scheme, support vector regression models were constructed using all applicable 2-dimensional molecular descriptors calculated by the Python package Mordred ("All Mordred MD"). In the second scheme, each model was tuned using recursive feature elimination (RFE) and utilized a varying subset of the Mordred descriptors ("Mordred MD + RFE"). The last selection scheme built models using only MD that were highly conserved (>95%) in the RFE trials for each ion-type; the number of MD used in $[M-H]^-$, $[M+H]^+$, and $[M+Na]^+$ trials were 75, 101 and 87, respectively ("Consensus MD"). The list of descriptors in each consensus set is available in the Supplemental Excel file. Each trial represents a prediction of 30% of a data set using the remaining 70% for training, and 101 predictions were performed for each combination of ion-type and MD selection scheme. Violin plots showing the distribution of the median relative percent error (a-c) and root mean square error (d-f) are presented.

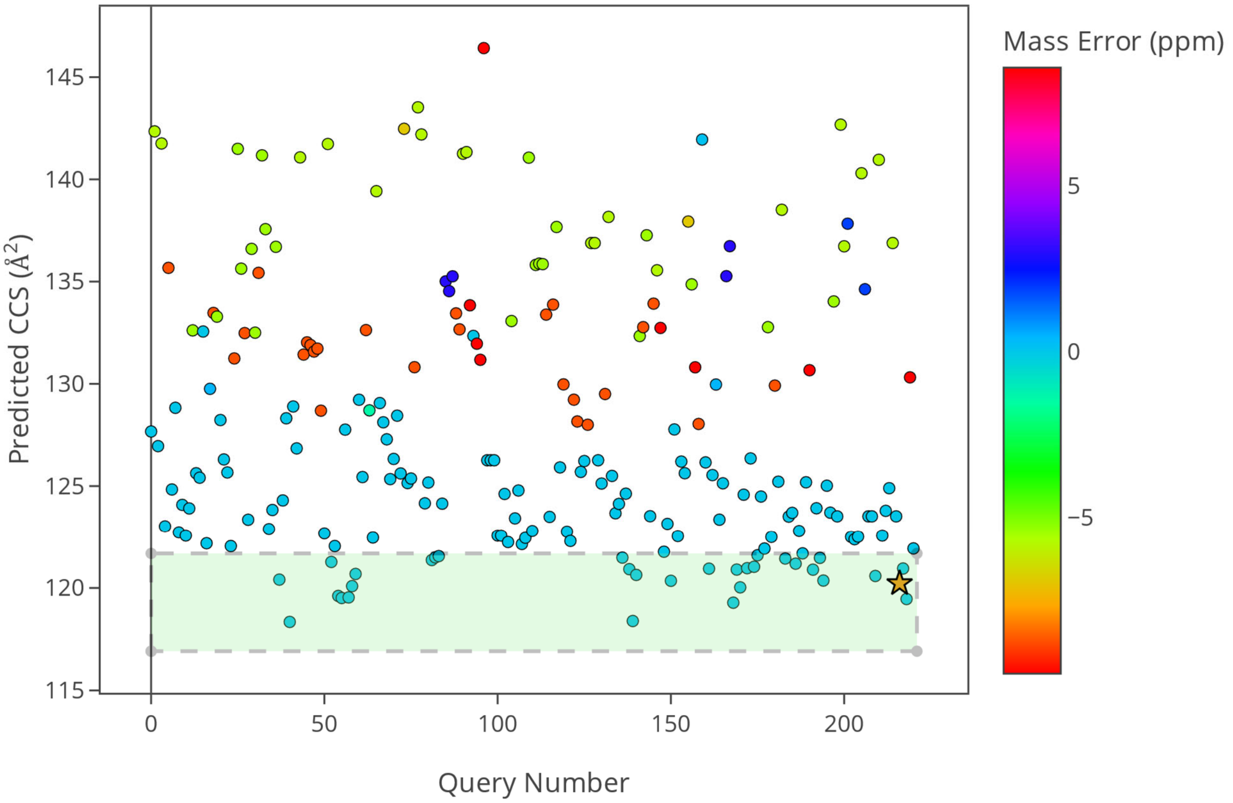


**Figure S8: Predicted CCS values for all PubChem candidates queried for the example presented in Figure 4**. A support vector regression model was trained to predict collision cross sections with a calibration set of singly deprotonated ions from the McLean Unified CCS Compendium ($n = 397$). Cross-validation using a 5-fold scheme yielded a root mean square error cross validation of 4.832 Å$^2$ and a median relative percent error of 1.221 %. The model was then tested on an external validation set from the Compendium ($n = 170$) and yielded a root mean square error validation of 5.222 Å$^2$ and a median relative percent error of 1.466 %. Green shaded region: experimental CCS value ± 2%; Gold star: predicted CCS value of isoorotic acid; Color scale: mass error (ppm) between the standard’s monoisotopic mass and the mass of the candidate.

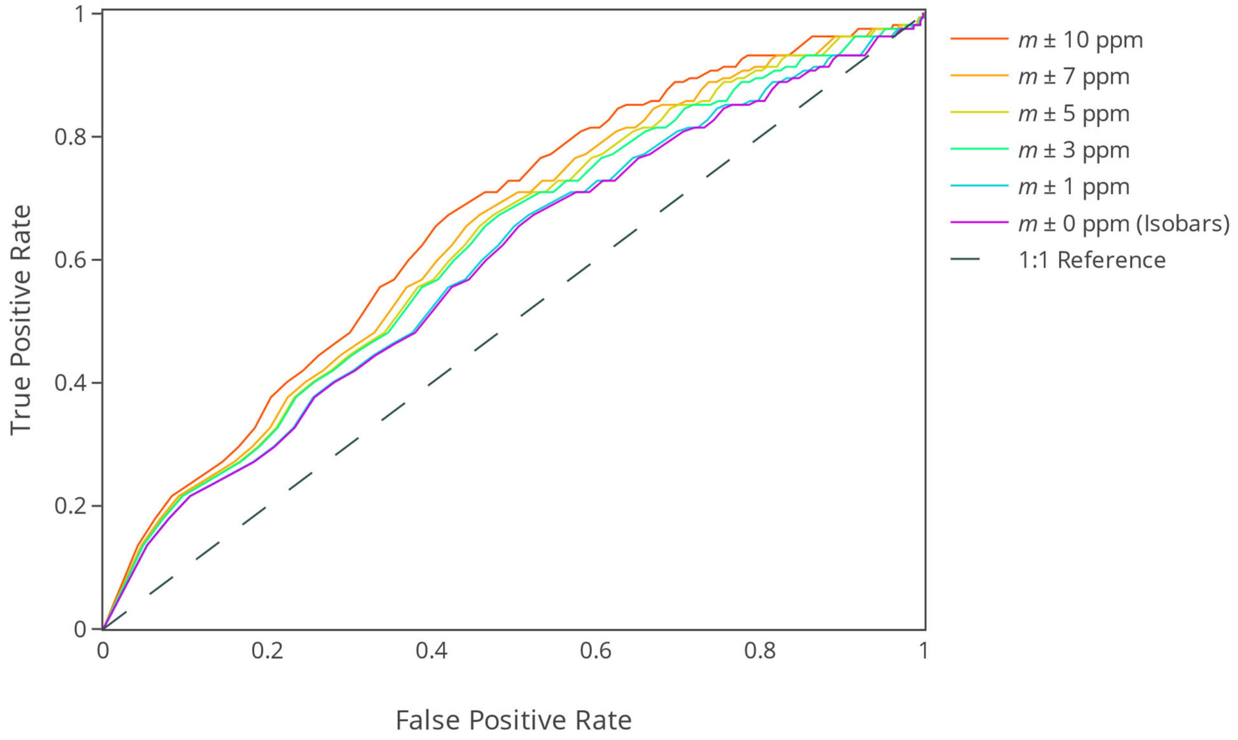


**Figure S9**: **Receiver Operating Characteristic (ROC) curves depicting true positive versus false positive rates for metabolite annotations using CCS prediction with the complete $[\text{M-H}]^-$ training and test sets**.

A data set of CCS values for $[\text{M-H}]^-$ ions was obtained from the McLean Unified CCS Compendium ($n = 559$). A total of 397 measurements were used to train support vector regression models to predict CCS values for new deprotonated species, while 162 were retained for ROC analysis. Each compound in the test set was treated as an unknown, with only its monoisotopic mass ($m$) and experimental CCS value being utilized. For each validation compound, all entries in PubChem with monoisotopic masses within 10 ppm of its monoisotopic mass were retrieved. Combined, the 162 validation compounds yielded 598,208 annotation candidates. The correct annotations for the 162 validation compounds were considered actual positives (true positives + false negatives), while the remaining 598,046 annotations were considered actual negatives (true negatives + false positives). CCSP 2.0 was used to predict the CCS value of each candidate, and the relative percent error between the candidate's predicted CCS and the corresponding experimental CCS was calculated. Various %CCS thresholds were investigated; correct annotations within the threshold were considered true positives (TP), while correct annotations outside of the threshold were considered false negatives (FN). Incorrect annotations within the threshold were considered false positives (FP), while incorrect annotations outside the threshold were considered true negatives (TN). The ROC curve represents true positive rate ($TPR = TP/(TP + FN)$ as a function of the false positive rate ($FPR = FP/(FP + TN)$). This process was repeated for annotation candidate lists generated with 7 ppm, 5 ppm, 3 ppm, 1 ppm, and 0 ppm mass windows. An ideal ROC curve would discover all correct candidates without discovering incorrect candidates, leading to an area under the curve (AUC) of 1. Equal true and false positive rates (grey dashed line) would show CCS cannot discern the correct annotation, leading to an AUC of 0.5.

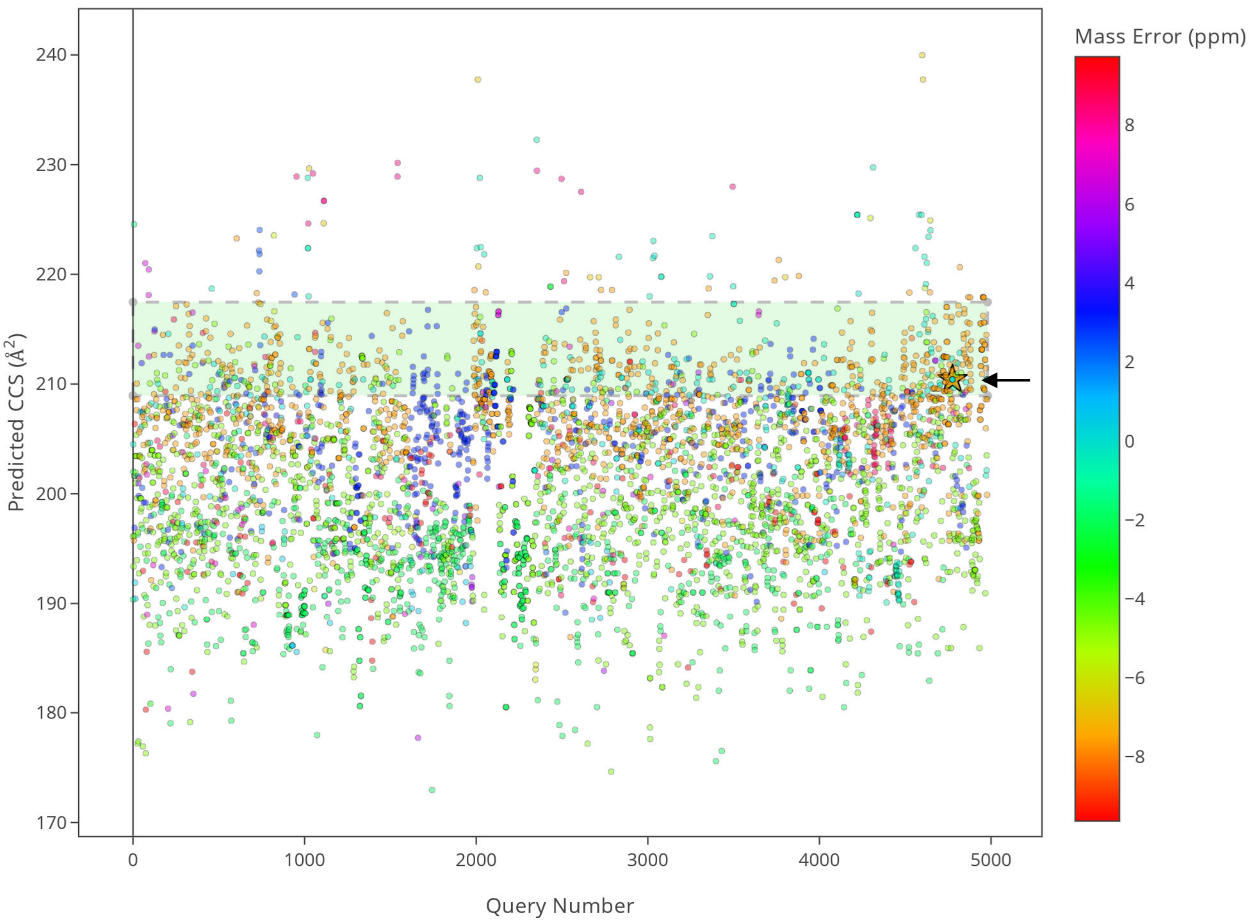


**Figure S10**: **Predicted CCS values for all PubChem candidates queried for the example presented in Figure 6**. A support vector regression model was trained to predict CCS values with a calibration set of $[M\text{-}H]^-$ ions obtained from members the lipid superclass in the McLean Unified CCS Compendium ($n = 93$). Cross-validation using a 5-fold scheme yielded a root mean square error cross validation of 5.268 Å$^2$ and a median relative percent error of 1.066 %. The model was then tested on an external validation set selected from the Compendium ($n = 36$), yielding a root mean square error validation of 3.463 Å$^2$ and a median relative percent error of 0.862 %. Green shaded region: experimental CCS value ± 2%; Gold star: predicted CCS value of isoorotic acid; Color scale: mass error (ppm) between the standard's monoisotopic mass and the mass of the candidate. Reduced marker opacity to improve visualization of overlapping points.

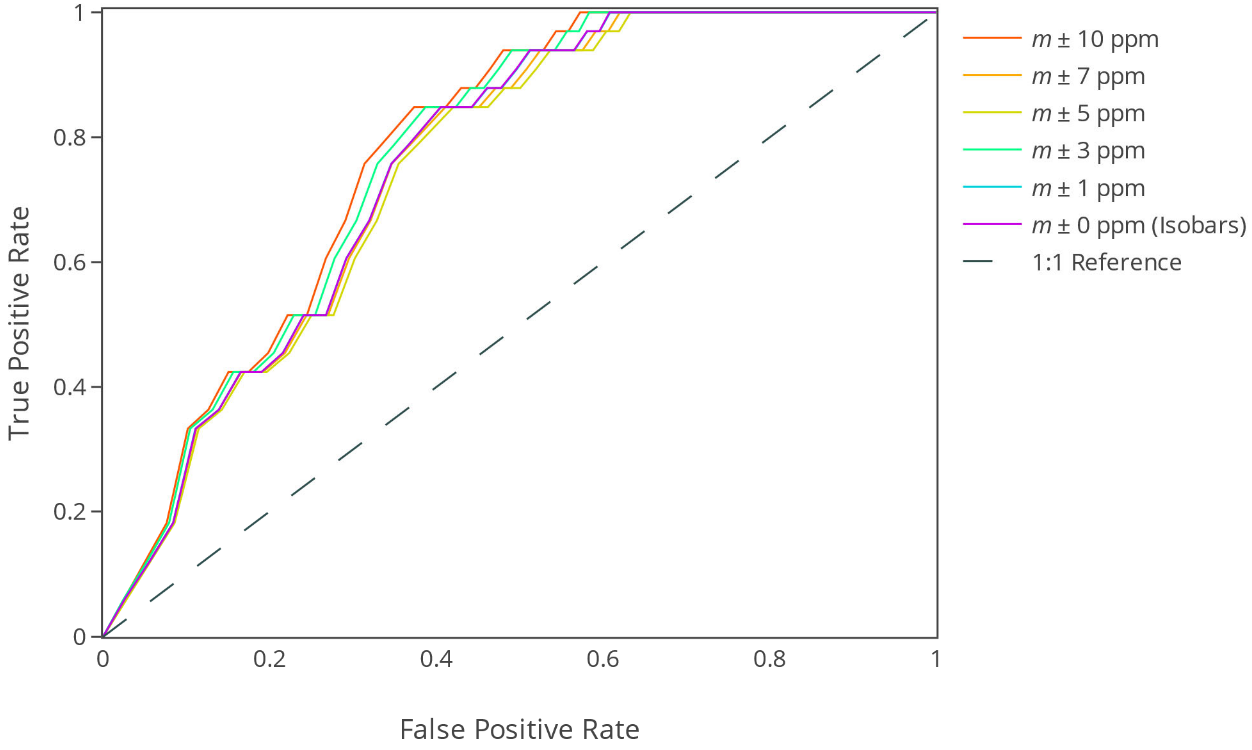


**Figure S11**: **Receiver Operating Characteristic (ROC) curves depicting true positive versus false positive rates for metabolite annotations using CCS predictions with $[\text{M-H}]^-$ lipid training and test sets**.

A set of $n = 126$ CCS values for $[\text{M-H}]^-$ ions was obtained from the lipid superclass of the McLean Unified CCS Compendium. The data set was randomly split with 93 being allocated to the training of SVR models, while 33 were retained for ROC analysis. Each compound in the test set was treated as an unknown, with only its monoisotopic mass ($m$) and experimental CCS value being utilized. For each validation compound, all entries in PubChem with monoisotopic masses within 10 ppm of its monoisotopic mass were retrieved. Combined, the 33 validation compounds yielded 136,940 annotation candidates. The correct annotations for the 33 validation compounds were considered actual positives, while the remaining 136,940 annotations were considered actual negatives. CCSP 2.0 was used to predict the CCS value of each candidate, and the relative percent error between the candidate's predicted CCS and the corresponding experimental CCS was calculated. Various %CCS thresholds were investigated; correct annotations within the threshold were considered true positives (TP), while correct annotations outside of the threshold were considered false negatives (FN). Incorrect annotations within the threshold were considered false positives (FP), while incorrect annotations outside the threshold were considered true negatives (TN). The ROC curve represents true positive rate ($TPR = TP/(TP + FN)$ as a function of the false positive rate ($FPR = FP/(FP + TN)$). This process was repeated for annotation candidate lists generated with 7 ppm, 5 ppm, 3 ppm, 1 ppm, and 0 ppm mass windows. An ideal ROC curve would discover all correct candidates without discovering incorrect candidates, leading to an area under the curve (AUC) of 1. Equal true and false positive rates (grey dashed line) would show CCS cannot discern the correct annotation, leading to an AUC of 0.5.

**Table S7**. Mordred descriptor modules, counts and basic meaning.

| Module number | Module name | Number of descriptors |
|---|---|---|
| 1 | ABCIndex | 2 |
| 2 | AcidBase | 2 |
| 3 | AdjacencyMatrix | 13 |
| 4 | Aromatic | 2 |
| 5 | AtomCount | 16 |
| 6 | Autocorrelation | 606 |
| 7 | BCUT | 24 |
| 8 | BalabanJ | 1 |
| 9 | BaryszMatrix | 104 |
| 10 | BertzCT | 1 |
| 11 | BondCount | 9 |
| 12 | CPSA | 2 |
| 13 | CarbonTypes | 10 |
| 14 | Chi | 56 |
| 15 | Constitutional | 16 |
| 16 | DetourMatrix | 14 |
| 17 | DistanceMatrix | 13 |
| 18 | EState | 316 |
| 19 | EccentricConnectivityIndex | 1 |
| 20 | ExtendedTopochemicalAtom | 45 |
| 21 | FragmentComplexity | 1 |
| 22 | Framework | 1 |
| 23 | HydrogenBond | 2 |
| 24 | InformationContent | 42 |
| 25 | KappaShapeIndex | 3 |
| 26 | Lipinski | 2 |
| 27 | LogS | 1 |
| 28 | McGowanVolume | 1 |
| 29 | MoeType | 53 |
| 30 | MolecularDistanceEdge | 19 |
| 31 | MolecularId | 12 |
| 32 | PathCount | 21 |
| 33 | Polarizability | 2 |
| 34 | RingCount | 138 |
| 35 | RotatableBond | 2 |
| 36 | SLogP | 2 |
| 37 | TopoPSA | 2 |
| 38 | TopologicalCharge | 21 |
| 39 | TopologicalIndex | 4 |
| 40 | VdwVolumeABC | 1 |
| 41 | VertexAdjacencyInformation | 1 |
| 42 | WalkCount | 21 |
| 43 | Weight | 2 |
| 44 | WienerIndex | 2 |
| 45 | ZagrebIndex | 4 |

1. Djoumbou Feunang, Y.; Eisner, R.; Knox, C.; Chepelev, L.; Hastings, J.; Owen, G.; Fahy, E.; Steinbeck, C.; Subramanian, S.; Bolton, E.; Greiner, R.; Wishart, D. S., ClassyFire: automated chemical classification with a comprehensive, computable taxonomy. *J Cheminform* **2016,** *8*, 61.
